# PDS5 proteins regulate the length of axial elements and telomere integrity during male mouse meiosis

**DOI:** 10.1101/763797

**Authors:** Alberto Viera, Inés Berenguer, Miguel Ruiz-Torres, Rocío Gómez, Andrea Guajardo, José Luis Barbero, Ana Losada, José A. Suja

## Abstract

Cohesin cofactors regulate the loading, maintenance and release of cohesin complexes from chromosomes during the mitotic cell cycle but little is known on their role during vertebrate meiosis. One such cofactor is PDS5, which exists in two versions in somatic and germline cells, PDS5A and PDS5B, with unclear functional specificity. Here we have analyzed their distribution and functions in mouse spermatocytes. We show that simultaneous elimination of PDS5A and PDS5B results in severe defects during prophase I while their individual depletion does not, suggesting a functional redundancy of the two factors. Shortened axial/lateral elements and a reduction of early recombination nodules are observed in the absence of both PDS5 proteins. Moreover, telomere integrity and their association to the nuclear envelope are severely compromised. As these defects occur without detectable reduction in chromosome-bound cohesin, we propose that the dynamic behavior of the complex, mediated by PDS5 proteins, is key for successful completion of meiotic prophase I.

## Introduction

Meiosis is a highly specialized cell division process that produces haploid gametes by performing two consecutive rounds of chromosome segregation after a single round of DNA replication of a diploid genome. The accurate segregation of chromosomes during meiosis relies on the proper achievement of the preceding processes of homologous chromosome pairing, synapsis and recombination (Bolcun-Filas and Handel, 2018). Meiotic recombination initiates in mammals during the leptotene stage of prophase I by the formation of programmed DNA double-strand breaks (DSBs) by the endonuclease SPO11 (Chicheportiche et al., 2007), which leads to phosphorylation of the histone variant H2AX at serine 139 (γ-H2AX) in the surrounding chromatin (Mahadevaiah et al., 2001). After a 5’ to 3’ resection of DNA at DSBs, the recombinase RAD51, among others, associates to the 3’ single-stranded DNA and promotes invasion of the double-stranded DNA of the homolog, a process that facilitates recognition and pairing of the homologs (Grabarz et al., 2012). Some of these events are finally processed as reciprocal crossovers via the participation of the DNA mismatch repair protein MLH1 (Moens et al., 2007). Although meiotic recombination initiates early in meiosis, its resolution takes place in the pachytene stage after the homologs have achieved synapsis. Synapsis is mediated by the synaptonemal complex (SC), a tripartite proteinaceous structure specific of meiosis that assembles during prophase I (Fraune et al., 2012). The SC assembly is initiated in leptotene, when an axial element (AE) is developed along each homolog over the trajectories of previously loaded cohesin complexes. During this stage, the telomeres attach to the nuclear envelope (NE), and by zygotene they adopt a polarized “bouquet” configuration that promotes chromosome movements essential for accurate homolog pairing (Shibuya and Watanabe, 2014). Subsequently, the so-called central element (CE) is formed between AEs of the SC. At this time, the AEs are referred as lateral elements (LEs) and the CE, composed by transverse filaments, physically connects the LEs of the two homologs. By pachytene, the homologs have achieved synapsis along their entire length and thus each bivalent presents a fully developed SC.

During the first and second meiotic divisions, recombined homologs and single chromatids migrate to opposite spindle poles, respectively, thanks to a sequential loss of arm and centromere sister-chromatid cohesion (Suja and Barbero, 2009; Ishiguro, 2019). Cohesion is mediated by ring-shaped cohesin complexes composed of two SMC (Structural Maintenance of Chromosomes) proteins, SMC1α and SMC3, the kleisin subunit RAD21 and a STAG/SA subunit that can be SA1 or SA2 in somatic vertebrate cells (Morales and Losada, 2018). The dynamic loading, maintenance and release of cohesin complexes from chromatin is mediated by cofactor proteins and some posttranslational modifications (Rankin, 2015; Ouyang and Yu, 2017). The functions of these cofactors have been characterized during the mitotic cycle, but much less is known of their behavior in meiosis. Once cohesin is loaded on chromatin in early G1 by the NIPBL-MAU2 heterodimer, PDS5 and WAPL bind to the complex and promote their dissociation (Tedeschi et al., 2013; Ouyang et al., 2016). During S phase, as sister chromatids arise from the replication fork, cohesion is established by a fraction of cohesin complexes that are acetylated in their SMC3 subunit and bound by Sororin, which counteracts the cohesin releasing activity of WAPL until mitotic prophase (Shintomi and Hirano, 2009; Nishiyama et al., 2010). Two versions of the PDS5 protein exist in vertebrate cells, PDS5A and PDS5B, which are required both for cohesin dissociation together with WAPL and for cohesin stabilization together with Sororin (Losada et al, 2005; Carretero et al., 2013; Ouyang et al, 2016; Wutz et al., 2017). While the two PDS5 proteins present these activities, centromeric cohesion defects are more apparent in *Pds5B* deficient cells in mitosis (Carretero et al., 2013).

In addition to the canonical cohesin subunits mentioned above, several meiosis-specific subunits have been described during mammalian meiosis. SMC1β and STAG3 are the paralogs of SMC1α and SA1/2, respectively, while REC8 and RAD21L are the meiotic counterparts of the kleisin subunit RAD21 (Ishiguro, 2019). The distribution and functions of the different cohesin subunits during mammalian meiosis has been extensively analyzed. All of them, both mitotic and meiosis-specific ones, are located at AEs/LEs during mouse prophase I. Spermatocytes from mouse deficient for the meiosis-specific cohesin subunits SMC1β (Revenkova et al., 2004; Novak et al., 2008; Adelfalk et al., 2009; Biswas et al., 2013), REC8 (Bannister et al., 2004; Xu et al., 2005), RAD21L (Herrán et al., 2011), and STAG3 (Fukuda et al., 2014; Hopkins et al., 2014; Llano et al., 2014; Winters et al., 2014), arrest at different stages of prophase I and display severe defects in the assembly and pairing of AEs/LEs, recombination and, in some cases, altered telomere structure. However, there are few studies describing the dynamics and involvement of cohesin cofactors during mammalian meiosis. WAPL decorates the AEs/LEs of chromosomes in mouse oocytes (Zhang et al., 2008) and spermatocytes, where it is involved in the removal of arm cohesion by the end of prophase I (Brieño-Enríquez et al., 2016). Unexpectedly, Sororin has been detected at the SC central region, unlike the cohesin subunits and WAPL (Gómez et al., 2016; Jordan et al., 2017).

The single orthologs of PDS5 in *Sordaria macrospora* (Spo76p) and budding yeast are located at AEs/LEs during prophase I stages (van Heemst et al., 1999; Zhang et al., 2005; Jin et al., 2009). Analyses of meiosis in Pds5 mutants in *Sordaria*, budding and fission yeasts, and the worm *Caenorhabditis elegans* indicate that this protein is involved in cohesion, formation of AEs/LEs, condensation, homolog pairing and repair of DSBs (van Heemst et al., 1999; Wang et al., 2003; Zhang et al., 2005; Ding et al., 2006; Jin et al., 2009; Ding et al., 2016). In mammaliam meiosis, like in mitosis, both PDS5A and PDS5B are present. While nothing is known about the localization of PDS5A, PDS5B has been detected at AEs/LEs in mouse spermatocytes (Fukuda and Höög, 2010). In the present study, we report the distribution of PDS5A and PDS5B during male mouse meiosis. Moreover, taking advantage of conditional knock out mouse models previously generated (Carretero et al., 2013), we have analyzed the meiotic phenotype of spermatocytes lacking one or both PDS5 proteins. Our results indicate that a single PDS5 protein, either PDS5A or PDS5B, is sufficient for prophase I progression. However, their simultaneous elimination results in drastic meiotic defects including the formation of shortened AEs/LEs, reduced formation of early recombination nodules, and alterations in the structure of telomeres and their attachment to the NE during prophase I.

## Results

### Different localization and dynamics of PDS5A and PDS5B in mouse spermatocytes

We analyzed the distribution of PDS5A and PDS5B proteins on spread mouse spermatocytes by immunofluorescence using specific antibodies for these proteins (Fig. S1A-C) and for SYCP3, a structural component of the AEs/LEs, to identify the different meiotic stages. PDS5A was undetectable during leptotene when SYCP3-labeled AEs started to organize (Fig. 1A and B), but colocalized with SYCP3 along the synapsing AEs/LEs of both autosomes and sex chromosomes during zygotene (Fig. 1C and D) and early pachytene (Fig. 1E and F). By mid pachytene, however, the PDS5A labeling dispersed throughout the chromatin (Fig. 1G) and was less intense on the sex body (Fig. 1H). In diplotene, lack of staining was observed around the ends of the desynapsing autosomal LEs which corresponded to DAPI-positive regions, namely the chromocenters (Fig. 1I-L). This distribution of PDS5A persisted in diakinesis although the overall signal intensity was slightly reduced (Fig. 1M and N). By metaphase I PDS5A was found at centromeres (Fig. 1O). In side-viewed centromeres, PDS5A was present as a T-shaped signal below the closely associated sister kinetochores stained with ACA (Fig. 1O; bottom inset), while viewed from the top PDS5A encircled sister kinetochores (Fig. 1O; top inset). This localization is similar to that of SYCP3 at the inner centromere domain (Parra et al., 2004; Gómez et al., 2007), but different from that of the meiosis-specific cohesin subunit REC8 which localizes in small patches along the interchromatid domain at the arms (Fig. 1Q and R). Both PDS5A and REC8 were detected at the inner centromere domain, but in side-viewed centromeres the PDS5A signals were larger than those of REC8 (Fig. 1Q, inset). In metaphase II PDS5A was also present at the centromeres. In side-viewed centromeres two PDS5A signals were detected below kinetochores (Fig. 1S; bottom inset), while in top-viewed centromeres a single PDS5A ring encircled kinetochores (Fig. 1S; top inset).

**Figure 1.**
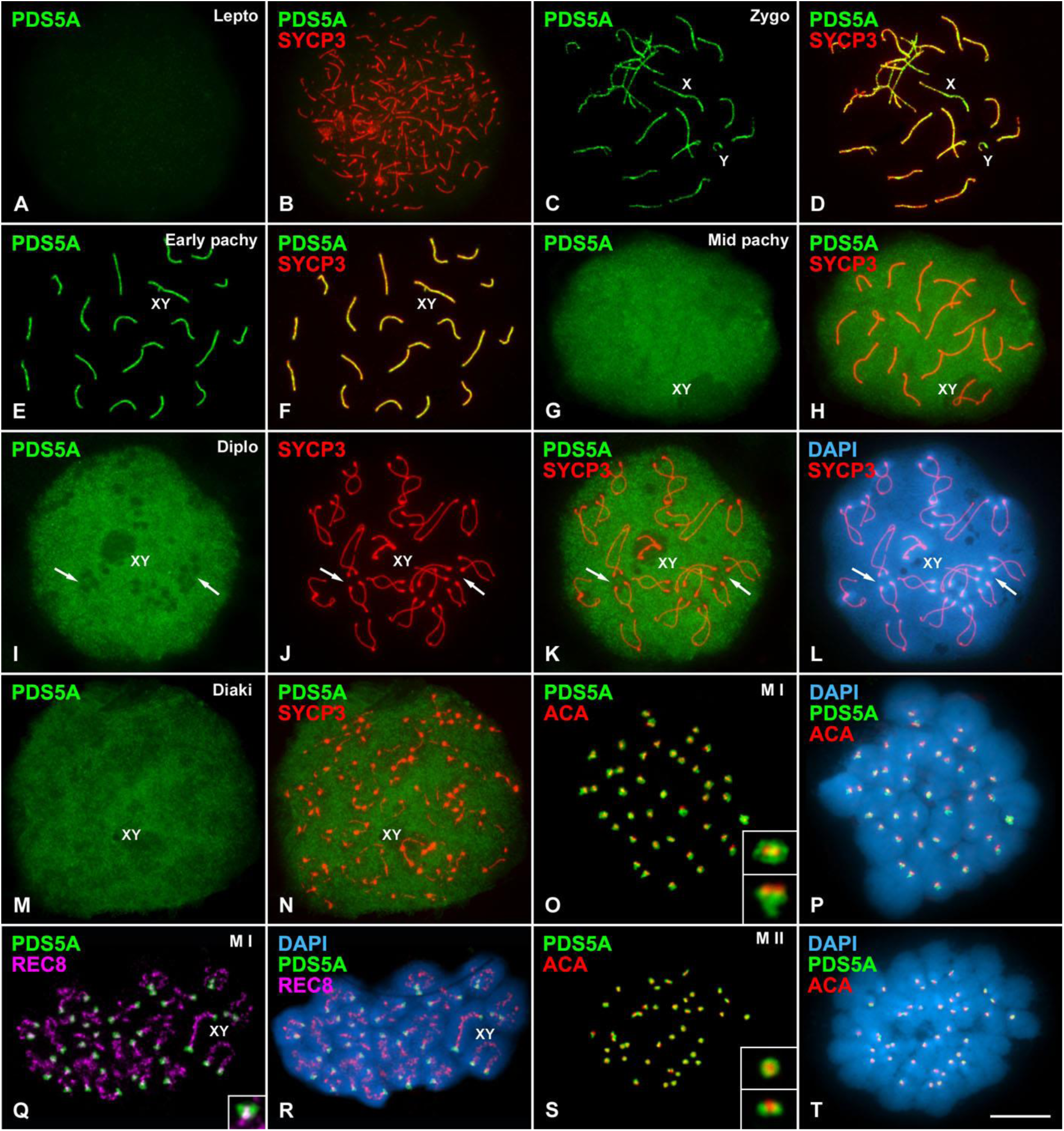
PDS5A distribution in mouse spermatocytes. **(A-N)** Double immunolabeling of PDS5A (green) and SYCP3 (red) in mouse spread spermatocytes in leptotene (A and B; Lepto), zygotene (C and D; Zygo), early pachytene (E and F; Early pachy), mid pachytene (G and H; Mid pachy), diplotene (I-L; Diplo) and diakinesis (M and N; Diaki). Arrows in I-L indicate the position of some chromocenters. **(O and P)** Double immunolabeling of PDS5A (green) and kinetochores revealed by an ACA serum (red) in a metaphase I spermatocyte (M I). **(Q and R)** Double immunolabeling of PDS5A (green) and REC8 (pseudocolored in purple) in a metaphase I spermatocyte (M I). **(S and T)** Double immunolabeling of PDS5A (green) and kinetochores (ACA, red) in a metaphase II spermatocyte (M II). The position of sex chromosomes (X, Y) and bivalents (XY) is indicated. DAPI staining of the chromatin (blue) is shown for some spermatocytes. Details included in O, Q and S correspond to a 300% magnification. Scale bar: 10 μm.

Unlike PDS5A, PDS5B colocalized with SYCP3 along the AEs/LEs of the autosomes and sex chromosomes from leptotene up to pachytene (Fig. 2A-F). In addition, rounded PDS5B signals protruded from the AEs/LEs ends in pachytene (Fig. 2E and F). REC8 colocalized with PDS5B along AEs/LEs, but was not present at the two close PDS5B signals at SC ends (Fig. 2G and H). Instead, PDS5B signals at AEs/LEs ends colocalized with TRF1, a telomere binding protein (Fig. 2I and J). We found that PDS5B was also enriched at the ends of aligned but non-synapsed AEs/LEs present in pachytene-like spermatocytes from mutants for the SC central element protein SYCE3 (*Syce3^−/−^*), and the transverse filament protein SYCP1 (*Sycp1^−/−^*) (Fig. S2). Thus, the presence of PDS5B at telomeres does not require synapsis. We also analyzed the distribution of PDS5B in spermatocytes from mice deficient for the meiosis-specific cohesin subunits SMC1β (Adelfalk et al., 2009) and REC8 (Biswas et al., 2016), which present variable degrees of telomere abnormalities. In *Smc1β^−/−^* pachytene-like spermatocytes, PDS5B localized along the shorthened SCs and unsynapsed AEs/LEs (Fig. S3A-C), and altered signals were present at SC ends (Fig. S3B and C), resembling those reported for telomeric DNA or telomere proteins in *Smc1β^−/−^* mouse spermatocytes (Adelfalk et al., 2009). In *Rec8^−/−^* spermatocytes PDS5B was also observed at shortened unsynapsed AEs (Fig. S3D-F). In this case, a low number of AEs ends presented a highly reduced PDS5B labelling (Fig. S3E and F), consistent with the reported existence of 3 to 4 chromosomes per spermatocyte presenting decreased levels of telomeric DNA and telomere proteins (Biswas et al., 2016). Altogether our results indicate that in mouse spermatocytes PDS5B specifically accumulates at telomeres during prophase I.

**Figure 2.**
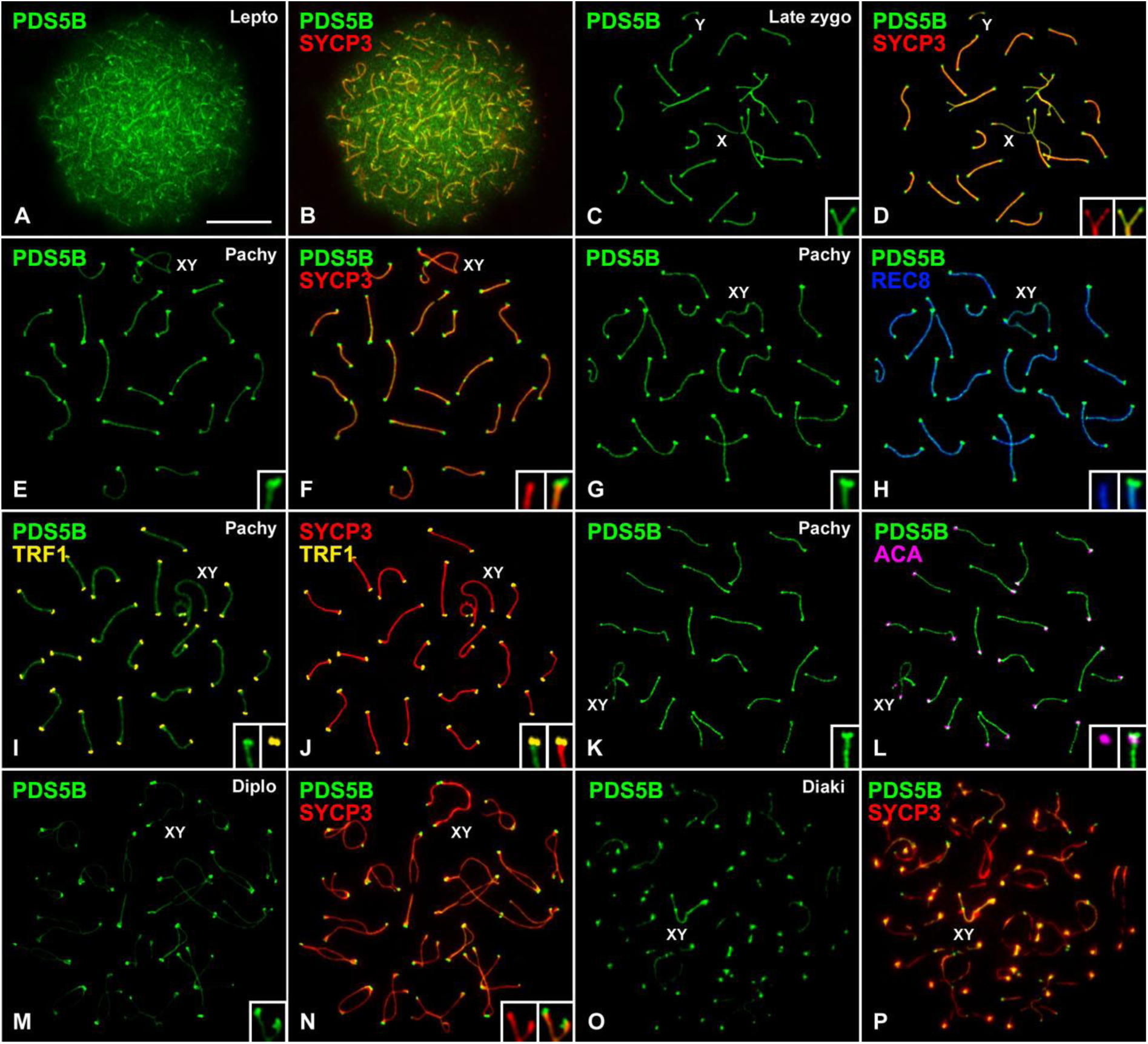
PDS5B distribution in prophase I spermatocytes. **(A-F)** Double immunolabeling of PDS5B (green) and SYCP3 (red) in spread spermatocytes in the stages indicated. **(G-L**) Immunolabeling of PDS5B (green) and either REC8 (pseudocolored in blue, G-H) or SYCP3 (red) and TRF1 (pseudocolored in yellow, I-J) or ACA (pseudocolored in purple, K-L) onto a spread pachytene spermatocyte (Pachy). **(M-P)** Double immunolabeling of PDS5B (green) and SYCP3 (red) on diplotene (M and N; Diplo) and diakinesis (O and P; Diaki) spread nuclei. The position of sex chromosomes (X, Y) and bivalents (XY) is indicated. Details included in C-P correspond to a 300% magnification. Scale bar: 10 μm.

Consistent with the telocentric nature of mouse chromosomes, kinetochore signals were always internally located in relation to PDS5B signals at the centromeric end of the pachytene LEs (Fig. 2K and L). By diplotene, PDS5B persisted along the desynapsing LEs, and at telomeres (Fig. 2M and N). In some images we could detect two closely associated PDS5B signals at desynapsed LEs (Fig. 2M and N, insets), corresponding to telomeres of sister chromatids. During diakinesis, PDS5B labeling began to disappear from the autosome LEs but not from telomeres (Fig. 2O and P). In metaphase I, large PDS5B signals colocalized with SYCP3 at the inner centromere domains (Fig. 3A-F). Interestingly, smaller PDS5B signals were located in pairs at the distal chromosome ends (Fig. 3C-F), that are reminiscent of the location of distal telomeres in metaphase I bivalents (Viera et al., 2003). In metaphase II chromosomes, large PDS5B signals were found at each centromere between the kinetochores (Fig. 3G-J). Altogether, our results indicate that PDS5A and PDS5B are present along the AEs/LEs in prophase I, similar to cohesin subunits, although the two paralogs display somehow different dynamics. Moreover, only PDS5B appears at telomeres.

**Figure 3.**
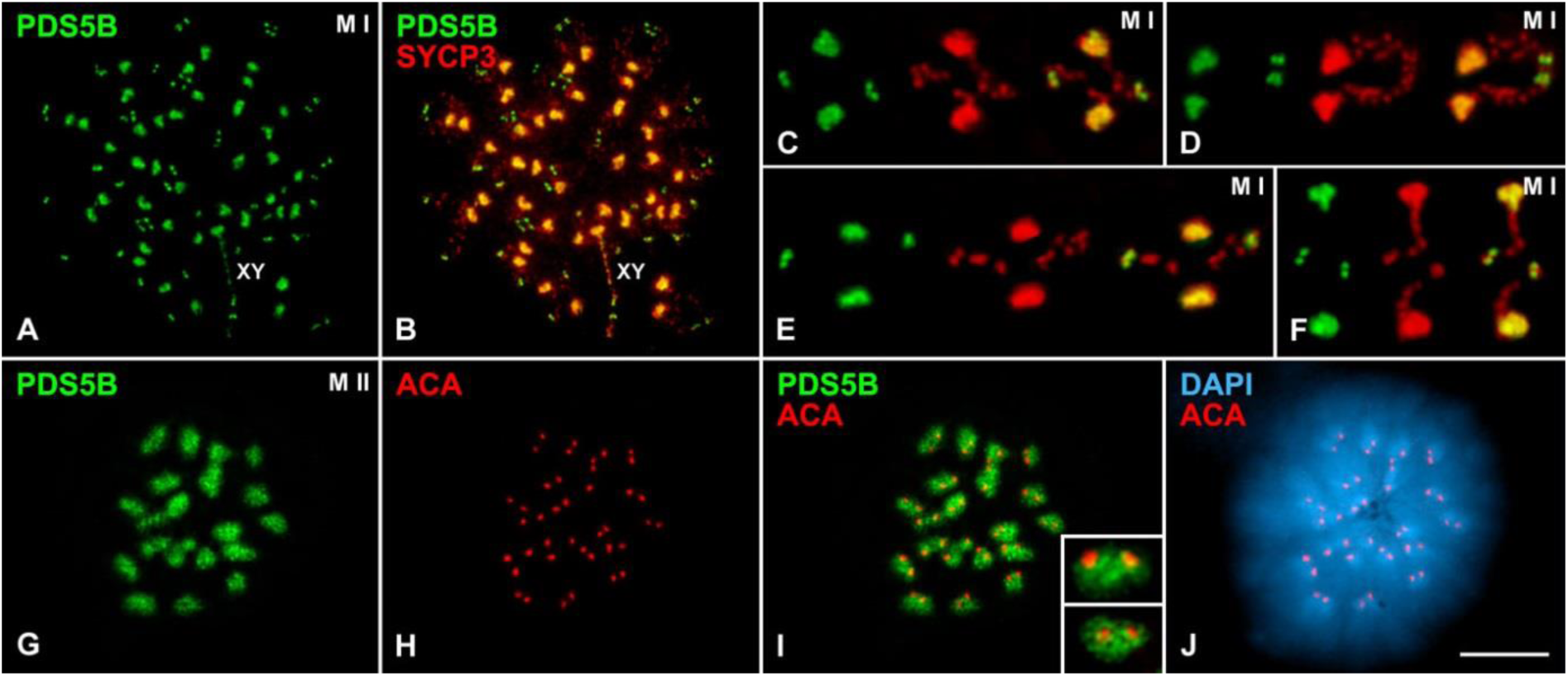
PDS5B distribution in meiotic condensed chromosomes. **(A-F)** Double immunolabeling of PDS5B (green) and SYCP3 (red) in metaphase I spermatocytes (M I). **(G-J)** Double immunolabeling of PDS5B (green) and kinetochores (ACA, red), and staining of the chromatin (DAPI) in a metaphase II spermatocyte (M II). The position of the sex bivalent (XY) is indicated. Details included in C-F and I correspond to a 300% magnification. Scale bar: 10 μm.

### A single PDS5 paralog protein is sufficient for prophase I progression

The different localization patterns of PDS5A and PDS5B during prophase I stages suggest that they may have specific functions in meiosis. To address this possibility, we took advantage of *Pds5* deficient mice generated previously (Carretero et al., 2013). Since constitutive knock out alleles for each gene are lethal in homozygosis, we used male mice carrying conditional knock out (cKO) alleles for *Pds5A*, *Pds5B* or both (*Pds5AB* hereafter) in combination with a Cre recombinase allele that can be induced by tamoxifen supplemented in the diet (Fig. S1A). Taking into account the approximate duration of meiosis in male mouse (Griswold, 2016), we initially designed an experimental procedure that could allow us to study the phenotype of *Pds5* deficient mice at all meiotic stages. We supplemented food pellets with 4-OHT to eight-week old wild-type, *Pds5A* cKO*, Pds5B* cKO and *Pds5AB* cKO mice for 6 weeks. Despite wild-type, *Pds5A* cKO and *Pds5B* cKO animals were healthy during the 6 weeks treatment, the health condition of double *Pds5AB* cKO individuals was negatively affected from day 15 onwards. Consequently in order to assure the animal’s welfare, and to standardize the experimental conditions, we decided to set the duration of 4-OHT treatments to 15 days, independently of the genotype. Efficient excision of the corresponding targeted exon was confirmed by PCR while immunoblot analyses of testis extracts showed only a partial decrease of protein levels (Fig. S1B and C). However, immunofluorescence analysis showed the absence of PDS5A and PDS5B proteins in tamoxifen-treated *Pds5A* cKO and *Pds5B* cKO spermatocytes, respectively, before mid-pachytene stage, while in later stages stainings were indistinguisable between wild-type and *Pds5* deficient cells (Fig. S1D-W). We reckon that the PDS5 proteins present in these mid-late pachytene spermatocytes were generated prior to the excision of the corresponding gene after initiation of the tamoxifen treatment. These results are congruent with the predicted duration of the different prophase I stages in mouse spermatogenesis (Griswold, 2016). Furthermore, the analysis of seminiferous tubules from both *Pds5A* cKO and *Pds5B* cKO individuals treated for 15-days with tamoxifen demonstrated that spermatocytes completed all meiotic and spermiogenic stages without any appreciable failure (Fig. S4).

Since PDS5B has been specifically involved in centromere cohesion in MEFs (Carretero et al., 2013), we next studied centromere distribution in *Pds5A* or *Pds5B* deficient spermatocytes. Our results did not evidence any appreciable alteration of centromere cohesion, or in their arrangement, in those spermatocytes (Fig. S5). Consequently, it can be assumed that neither PDS5A nor PDS5B are essential for prophase I progression until mid pachytene in male mice. However, simultaneous ablation of the two genes does have an important impact, as described in the next sections.

### PDS5 proteins regulate AE/LE length but are not required for synapsis

We next analyzed spermatocytes from *Pds5AB* cKO animals after tamoxifen treatment. Both PDS5A and PDS5B were completely absent from leptotene, zygotene and some pachytene spermatocytes (Fig. 4A-C and F-H). Interestingly, these early prophase I spermatocytes presented short SYCP3-labeled AEs/LEs and SCs (Figs. 4A-C and F-H, and S6A-C), a phenotype characteristic of all meiotic cohesin subunit mutants (Biswas et al., 2016; Ward et al., 2016). In some other pachytene spermatocytes wild-type levels of PDS5A and PDS5B were present and showed normal length SCs (Figs. 4D and I, and S6D). Metaphase I bivalents also showed wild-type levels and distributions of PDS5A and PDS5B (Fig. 4E and J), and meiotic divisions and spermiogenesis looked normal (Fig. S6E-L and N-O). Staining with an antibody against the testis-specific histone H1t, a marker of mid-pachytene chromatin (Drabent et al., 1996), further showed that 95% of pachytene spermatocytes from *Pds5AB* cKO treated animals with short SCs (n=200) were H1t-negative (Fig. 4K), while the remaining 5% of cells were faintly positive for H1t labeling (Fig. 4L), and all pachytene spermatocytes with normal SC length were strongly positive for H1t (Fig. 4M). Thus, spermatocytes with short AEs/LEs are probably those generated during tamoxifen treatment, which had progressed until early pachytene, while spermatocytes from mid pachytene onwards correspond to those that entered meiosis prior to the treatment. Average SC length measured in early pachytene spermatocytes (negative for H1t staining) was significantly reduced in *Pds5AB* deficient cells compared to wild-type (80±7 µm versus 154±20 µm, Fig. 4N), representing a reduction of around 52% of SC length. We also scored the number of spermatocytes in the different phases of prophase I (n=1,000) and observed a >2-fold increase in zygotene and a decrease in pachytene in *Pds5AB* deficient testes compared to wild-type (Fig. S6M and Q). Among the *Pds5AB* cKO pachytene spermatocytes only 11.6% of them displayed shortened SCs. Altogether, these results reflect a delay in synapsis and/or meiotic recombination progression.

**Figure 4.**
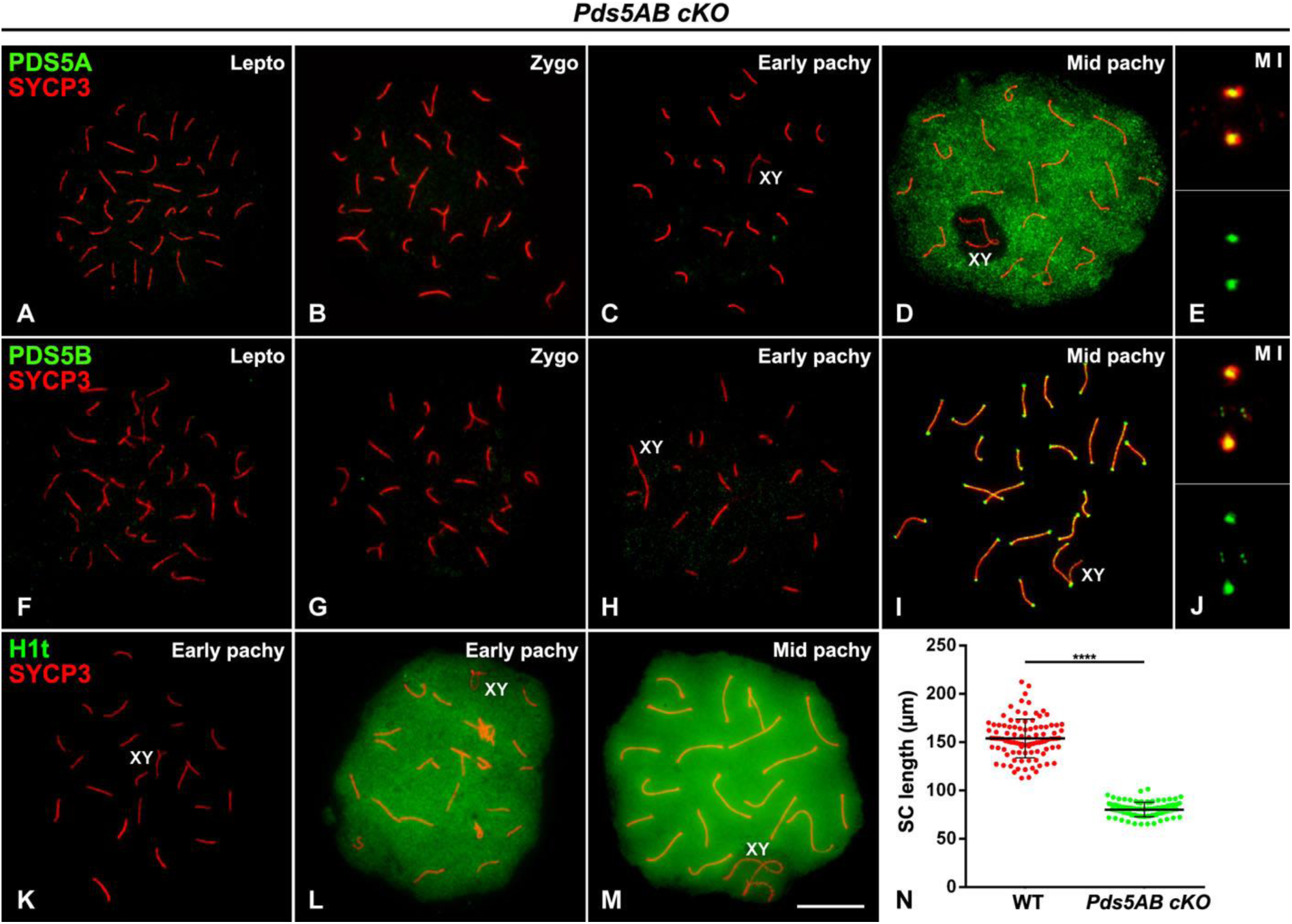
Reduced SC length in spermatocytes lacking PDS5A and PDS5B. **(A-E)** Double immunolabeling of PDS5A (green) and SYCP3 (red) in spread spermatocytes from tamoxifen-treated *Pds5AB* cKO in the stages indicated. **(F-J)** Similar stainings for PDS5B. **(K-M)** Preparations from the same mice were immunolabeled with the testis-specific histone variant H1t (green) and SYCP3 (red). The position of sex bivalents (XY) is indicated. Scale bar: 10 μm. **(N)** Scatter dot-plot graph of total synaptonemal complex (SC) length measured in the autosomal pairs of 33 early pachytene cells from three different individuals of each genotype. Bars indicate mean and standard error. Statistical significance was assessed using a T-test function (****, p < 0.0001).

To assess synapsis, we used double immunolabeling of SYCP3 and SYCP1. Leptotene wild-type spermatocytes displayed long discontinuous threads of SYCP3 representing the assembly of AEs along each chromosome (Fig. 5A). By contrast, 40 short individual SYCP3-labeled AEs were found in *Pds5AB* deficient leptotene spermatocytes (Fig. 5E). The pair-wise association of some AEs signals the initiation of synapsis by early zygotene (Fig. 5B and F). In these spermatocytes, the SYCP1 labeling was only located along the synapsed regions of the AEs/LEs (Fig. 5B and F), which became longer as synapsis progressed (Fig. 5C and G). Both the SYCP3-labeled AEs/LEs and the SYCP1-labeled CE were considerably shorter in the absence of both PDS5 proteins (Fig. 5C and G), but by early pachytene 19 fully synapsed autosomal SCs and the partially unsynapsed sex bivalent were found both in wild-type and *Pds5AB* deficient cells (Fig. 5D and H). Double labeling of SYCP3 and HORMAD2, a protein present at unsynapsed AEs/LEs (Wojtasz et al., 2009), further confirmed the accurate synapsis in *Pds5AB* deficient spermatocytes (Fig. 5I-L). We therefore conclude that PDS5 proteins are not required for synapsis of homologous chromosomes in male mouse meiosis.

**Figure 5.**
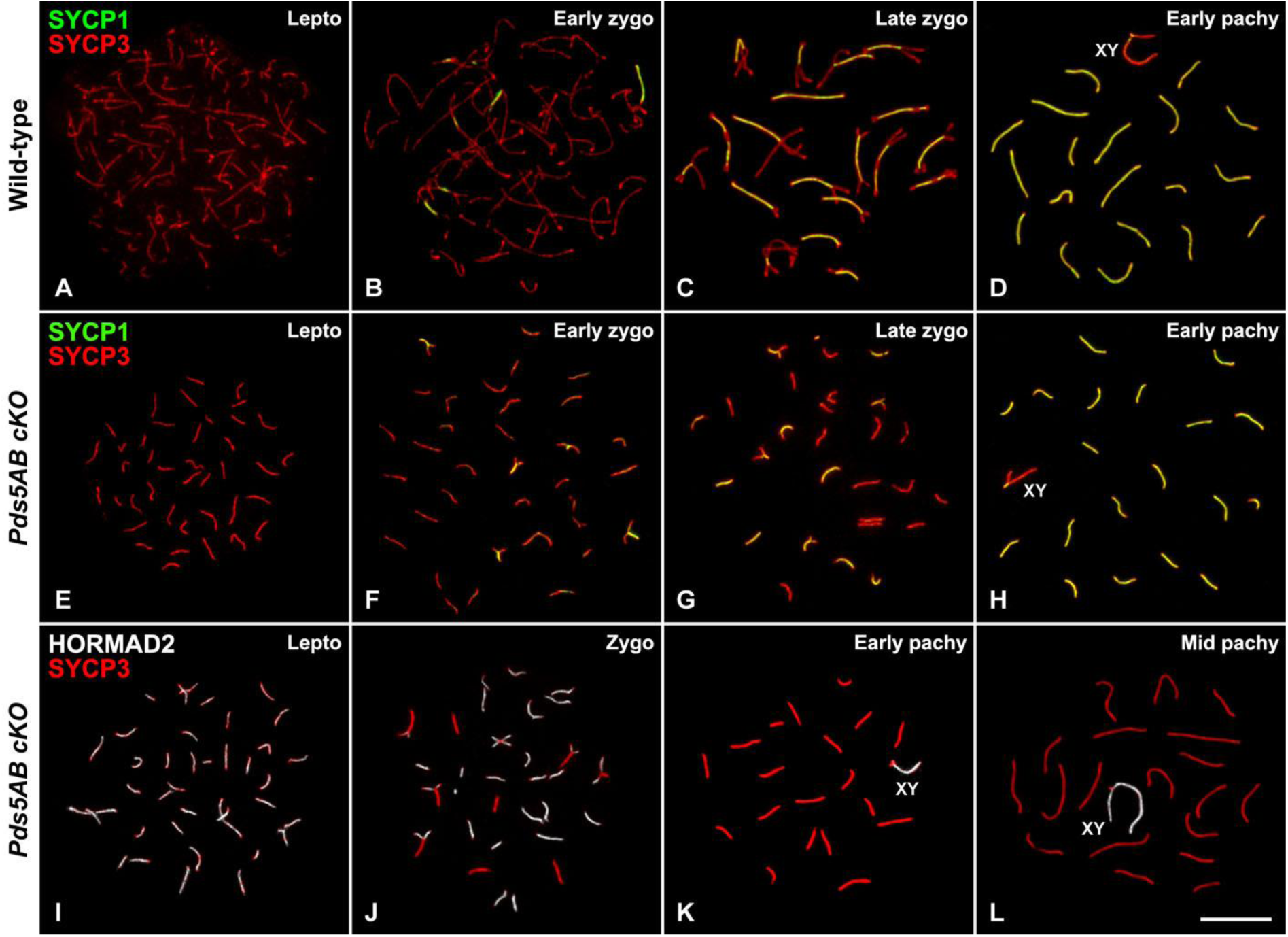
Synapsis progression in double *Pds5AB* cKO spermatocytes. **(A-H)** Double immunolabeling of SYCP1 (green) and SYCP3 (red) in spread spermatocytes from wild-type (A-D) and tamoxifen-treated *Pds5AB* cKO mice (E-H) in the stages indicated. **(I-L)** Double immunolabeling of HORMAD2 (pseudocolored in white) and SYCP3 (red).The position of sex bivalents (XY) is indicated. Scale bar: 10 μm.

### Normal cohesin distribution along meiotic chromosome axes in the absence of PDS5 proteins

We next analyzed whether the shortening of AEs/LEs in *Pds5AB* deficient spermatocytes was the consequence of an abnormal distribution or absence of cohesin complexes. Contrary to this possibility, the cohesin subunits SMC1α, SMC1β, SMC3, RAD21, RAD21L, REC8, and STAG3, were detected along the shorthened AEs/LEs in zygotene and early pachytene spermatocytes (Figs. 6A-I, and S7A-L). Cohesin cofactors Sororin and WAPL were also found along the synapsed regions (Fig. 6J-L) and AEs/LEs (Fig. 6M-O), respectively, as previously reported in wild-type spermatocytes (Brieño-Enríquez et al., 2016; Gómez et al., 2016; Jordan et al., 2017). Centromeric cohesion defects were not evident in deficient *Pds5AB* zygotene and early pachytene spermatocytes, as kinetochore signals appeared unaltered at the ends of AEs/LEs or SCs, respectively, as in *Pds5AB* proficient mid pachytene spermatocytes (Fig. S7M-O). Thus, the absence of PDS5 proteins does not compromise the loading and maintenance of cohesin complexes or centromere cohesion in prophase I.

**Figure 6.**
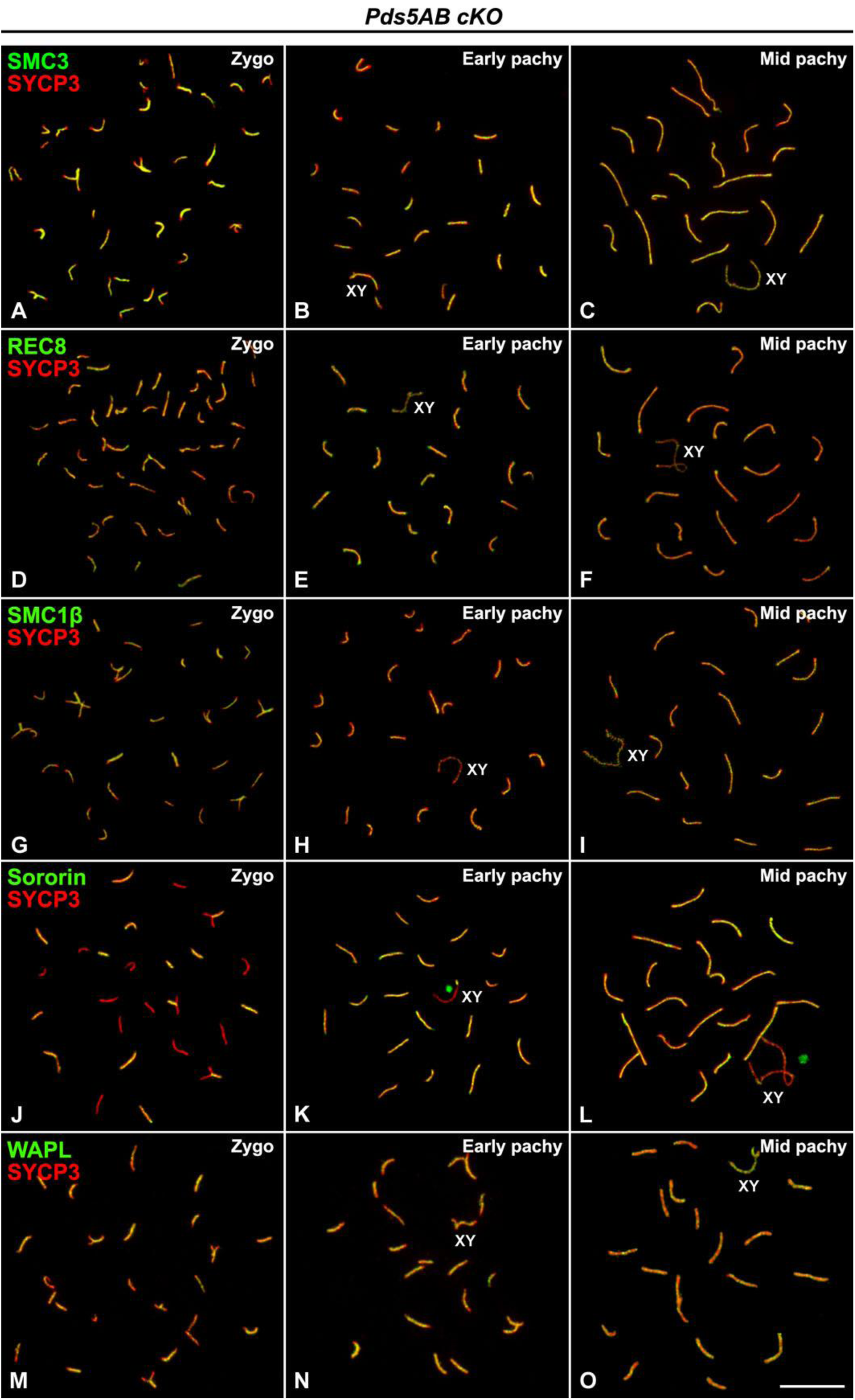
Distribution of cohesin subunits and regulators is not altered in the absence of PDS5A and PDS5B. **(A-O)** Double immunolabeling of SYCP3 (red) and the cohesin subunits SMC3 (green in A-C), REC8 (green in D-F) and SMC1β (green in G-I) and cohesin cofactors Sororin (green in J-L) and WAPL (green in M-O) in spread spermatocytes from tamoxifen-treated *Pds5AB* cKO mice in the stages indicated. The position of sex bivalents (XY) is indicated. Scale bar: 10 μm.

### PDS5 proteins affect meiotic recombination

Since *Pds5AB* deficient spermatocytes accumulate at zygotene but no clear defects in synapsis could be detected, we next asked about progression of meiotic recombination. The DSB marker γ-H2AX was found throughout the chromatin at leptotene, in chromatin surrounding unsynapsed regions in zygotene, and restricted to the sex body in early pachytene spermatocytes with shortened SCs, as well as in mid/late pachytene spermatocytes displaying a regular SC length (Fig. 7A-D). This dynamic distribution of γ-H2AX in *Pds5AB* deficient spermatocytes is consistent with that reported for wild-type spermatocytes (Mahadevaiah et al., 2001), suggesting that DSBs were generated and repaired in the absence of PDS5 proteins.

**Figure 7.**
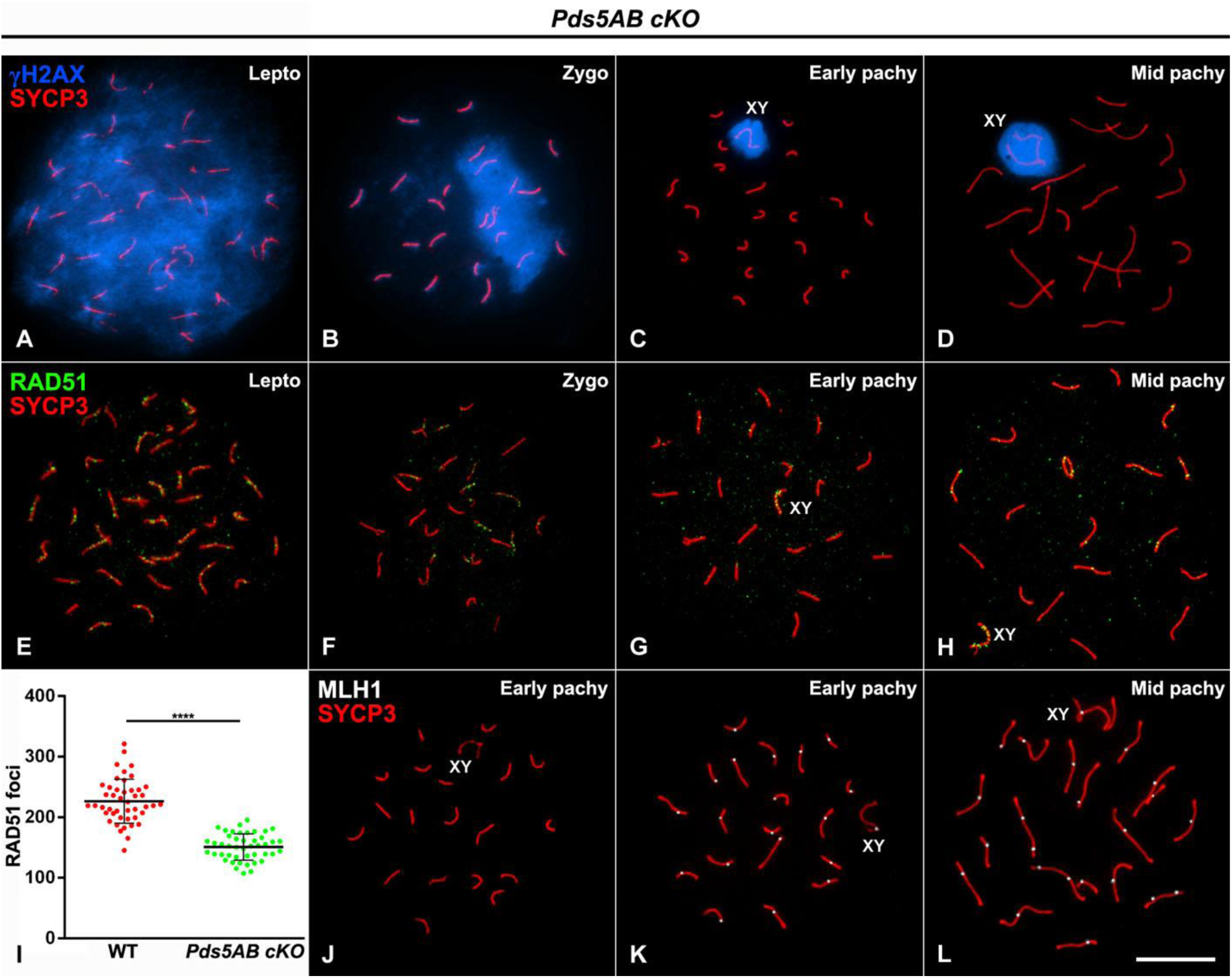
PDS5 proteins affect meiotic recombination. **(A-H)** Double immunolabeling of SYCP3 (red) and either γH2AX (pseudocolored in blue, A-D) or RAD51 (green, E-H) in spread spermatocytes from tamoxifen-treated *Pds5AB* cKO mice in the stages indicated. **(I)** Scatter dot-plot graph of the quantification of RAD51 foci in in 15 zygotene nuclei randomly selected from three different individuals of each genotype. Bars indicate mean and standard error and statistical significance was assessed with a T-test function (****, p < 0.0001). **(J-L)** Double immunolabeling of MLH1 (pseudocolored in white) and SYCP3 (red) in spread spermatocytes from tamoxifen-treated *Pds5AB* cKO mice. The position of sex bivalents (XY) is indicated. Scale bar: 10 μm.

Next, we analyzed the distribution of RAD51, a protein that participates in homology search at the so-called early recombination nodules along leptotene and zygotene AEs/LEs (Moens et al., 2007). RAD51 signals were indeed seen along AEs in leptotene spermatocytes with or without PDS5 proteins and their number gradually decreased as prophase I progressed (Figs. 7E-H, and S8). The average number of RAD51 foci was 226±36 in wild-type but only 151±22 in *Pds5AB* deficient zygotene spermatocytes (Fig. 7I). MLH1 foci, which mark late recombination nodules from mid-pachytene onwards (Moens et al., 2007), could not be detected in most *Pds5AB* deficient early pachytene spermatocytes with short SCs (96 out of 100; Fig. 7J). In the remaining 4 spermatocytes a single focus was invariably found in autosomal bivalents and in the sex bivalent (Fig. 7K), and therefore the mean number of MLH1 foci was 20. In mid/late pachytene spermatocytes displaying normal length SCs, the average number of MLH1 foci was 22±1 (n=91). Altogether, our results indicate that PDS5 proteins are not essential for initiation of meiotic recombination, but their absence reduces the number of early recombination nodules.

### Severe telomere abnormalities in the absence of PDS5 proteins

Since PDS5A and PDS5B participate in telomere cohesion in somatic cells (Carretero et al., 2013), and PDS5B is present at telomeres of meiotic chromosomes (Fig. 2), we analyzed the morphology and behavior of telomeres in *Pds5AB* deficient spermatocytes. Labeling with antibodies against the telomere proteins TRF1 (Fig. 8A-D) and RAP1 (Fig. 8E-H) evidenced the existence of multiple morphological alterations in early prophase I stages that were not found in wild-type, *Pds5A* deficient or *Pds5B* deficient spermatocytes. These alterations included stretched signals, multiple signals, a signal separated from the AEs/LEs, and ends without signals (i to iv in Fig. 8I and K). Moreover, they were similar for TRF1 and RAP1 staining, and could be detected in leptotene, zygotene and early pachytene spermatocytes (Fig. 8A-C and E-G, quantification in Fig. 8J and L, Table S1), but not in mid/late pachytene spermatocytes in which PDS5 levels are presumably normal (Fig. 8D and H). As a rule, more than 30% of telomeres presented an altered arrangement in zygotene and early pachytene *Pds5AB* spermatocytes (Fig. 8J and L, Table S1). We also studied the distribution of SUN1, an inner nuclear membrane protein that participates in the attachment of telomeres to the NE during prophase I, a crucial event for homologs pairing, recombination and synapsis in mammals (Ding et al., 2007). SUN1 signals appeared at some, but not all, AEs/LEs ends during leptotene, zygotene and early pachytene (Fig. 8M-O), and displayed altered morphologies reminiscent of the aberrations observed with TRF1 and RAP1 stainings (Fig. 8Q and R; quantification in Fig. 8S and Table S1). Anomalous or absent SUN1 signals were observed at 36.51% and 19.27% of telomeres in *Pds5AB* zygotene and early pachytene spermatocytes, respectively (Fig. 8M-S; Table S1). By mid/late pachytene, spermatocytes presented normal SUN1 signals at both ends of the LEs (Fig. 8P).

**Figure 8.**
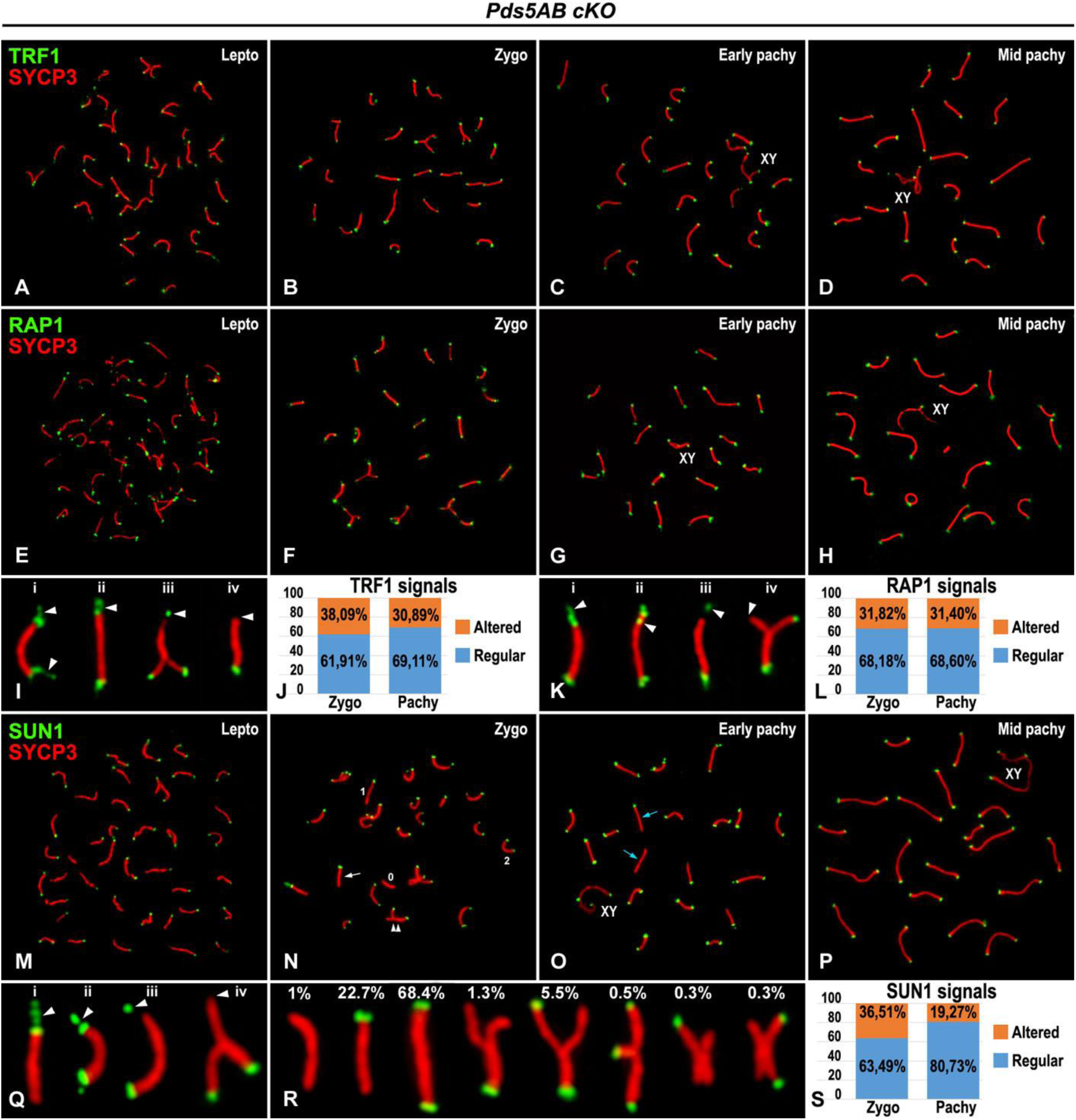
Altered distribution of telomeric proteins in the absence of PDS5 proteins. **(A-H)** Double immunolabeling of SYCP3 (red) and either TRF1 (green, A-D) or RAP1 (green, E-H) in spread spermatocytes from tamoxifen-treated *Pds5AB* cKO mice in the stages indicated. **(I)** Selected zygotene and pachytene bivalents presenting altered TRF1 distribution as (from left to right) (i) telomere stretches, (ii) multiple telomere, (iii) distant telomeres and (iv) telomere-less. **(J)** Quantification of telomeres with a regular or altered disposition of TRF1 in spermatocytes from the stages indicated lacking both PDS5 proteins. **(K, L)** As in I, J, respectively, but for RAP1 staining. **(M-P)** Double immunolabeling of SUN1 (green) and SYCP3 (red) in spermatocyte spreads from tamoxifen-treated *Pds5AB* cKO animals in the stages indicated. **(Q)** As in I, but for SUN1 staining. **(R)** Selected zygotene and pachytene bivalents from *Pds5AB* deficient spermatocytes immunolabeled with SYCP3 (red) and SUN1 (green), and their respective percentage of appearance. **(S)** Quantification of telomeres with a regular or altered disposition of SUN1. For further details on quantification of telomere aberrations see Table S1. The position of sex bivalents (XY) is indicated. Scale bar: 10 μm.

Fluorescence in situ hybridization (FISH) with a telomere-specific probe further confirmed the abnormal telomere organization in leptotene, zygotene and early pachytene spermatocytes lacking the two PDS5 proteins (Fig. 9A-D, quantification in 9E). By contrast, from mid/late pachytene onwards the telomere DNA showed a regular distribution at LEs ends (Fig. 9U-X), as in prophase I stages from *PdsA* deficient and *Pds5B* deficient spermatocytes (Fig. S9). A combined analysis of telomeric DNA by FISH and immunolabeling of RAP1 and SYCP3 in *Pds5AB* deficient spermatocytes in leptotene (Fig. 9F-J), zygotene (Fig. 9K-O) and early pachytene (Fig. 9P-T) demonstrated that a given telomere abnormality could be sometimes equally distinguished by FISH or immunolabeling (Fig. 9I and N), while in some other cases abnormal telomere FISH signals and no RAP1 signals were observed (Fig. 9J, O, S, and T). From these data we conclude that the absence of both PDS5 proteins promote the appearance of severe abnormalities at telomeres including aberrant telomeric DNA organization and distribution of telomere proteins.

**Figure 9.**
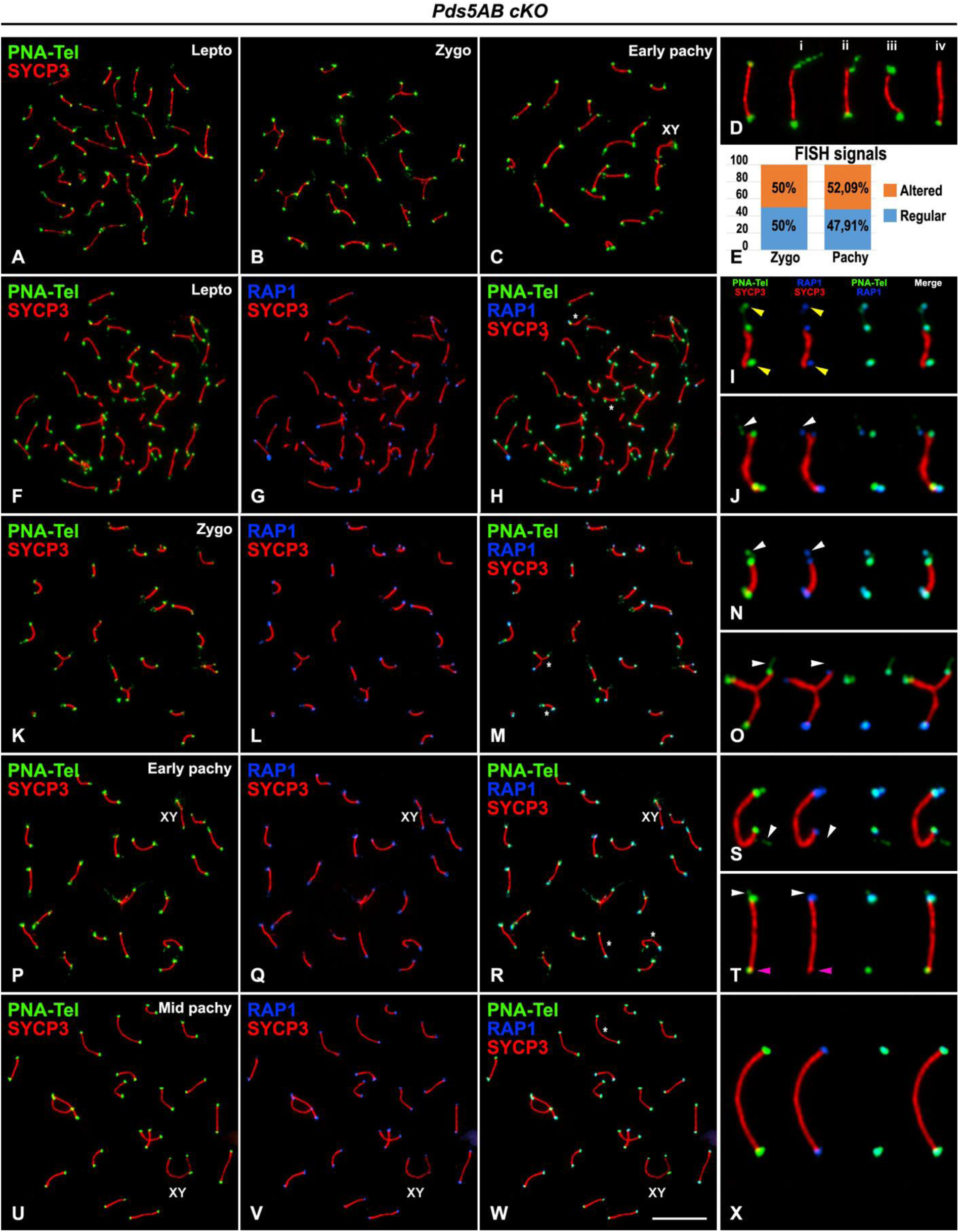
Altered organization of telomeric DNA in the absence of PDS5 proteins. **(A-C)** Immunolabeling of SYCP3 (red) and telomere FISH (PNA-Tel, green) in spread spermatocytes from tamoxifen-treated *Pds5AB* cKO mice. **(D)** 300% magnification of selected zygotene and pachytene bivalents presenting regular (left) or altered telomere FISH signals. **(E)** Quantification of telomere aberrations. **(F-X)** Double immunolabeling of SYCP3 (red) and RAP1 (blue) and telomere FISH (PNA-Tel, green) in spermatocytes from tamoxifen-treated *Pds5AB* cKO mice. Asterisks on H, M, R and W indicate the position of the enlarged chromosomes/bivalents shown in a 300% magnification in I, J, N, O, S, T, and X, respectively, displaying telomere stretches, multiple telomere, distant telomeres and telomere-less. Yellow arrowheads in I indicate telomeres displaying a distant telomere configuration observed both by telomere FISH and RAP1 immunolabeling. White arrowheads in J, N, O, S and T indicate telomeres with an altered organization of telomeric DNA into multiple telomere signals with (N) or without RAP1 signals (J, and T), and telomere stretches (O and S) without RAP1 signals. Pink arrowheads in T indicate a telomere presenting a regular telomere FISH signal without a RAP1 signal. The position of sex bivalents (XY) is indicated. Scale bar: 10 μm.

### PDS5 proteins promote proper attachment of telomeres to the NE

Given the alterations found at telomeres in *Pds5AB* deficient spermatocytes, we subsequently asked whether telomere attachment to the NE might be affected. To explore this possibility, we initially performed a double immunolabeling of SYCP3 and TRF1 on squashed spermatocytes, a technique that preserves the nuclear volume and chromosome positioning inside nuclei (Page et al., 1998; Parra et al., 2002), and collected image stacks across the entire volume of prophase I nuclei (Fig. 10). TRF1 signals were found at the ends of the SYCP3-labeled LEs in zygotene and early pachytene spermatocytes, from both wild-type and *Pds5AB* deficient mice (Fig. 10A-P). Nonetheless, while TRF1 signals were invariably positioned at the nuclear periphery in wild-type spermatocytes (Fig. 10A-D and I-L), some TRF1 signals were instead found at the nuclear interior in the absence of PDS5 proteins, indicating a lack of attachment of some telomeres to the NE (Fig. 10E-H and M-P; Movie S1). Morphological alterations in telomere structure similar to those observed in spread spermatocytes could be also detected (Fig. 10Q-T; Movie S2-S5). Interestingly, telomere-less ends of LEs were always found at the nuclear interior and thus non-attached to the NE (Movie S5). Similar results were obtained when analysing SUN1 distribution in squashed spermatocytes (Fig. S10; Movies S6-S8). As previously described for TRF1, SUN1 telomere-less ends of LEs were always located at the nuclear interior (Movie S8). Consequently, our data indicate that the simultaneous absence of PDS5A and PDS5B causes severe abnormalities in telomere morphology in prophase I spermatocytes and alters their attachment to the NE.

**Figure 10.**
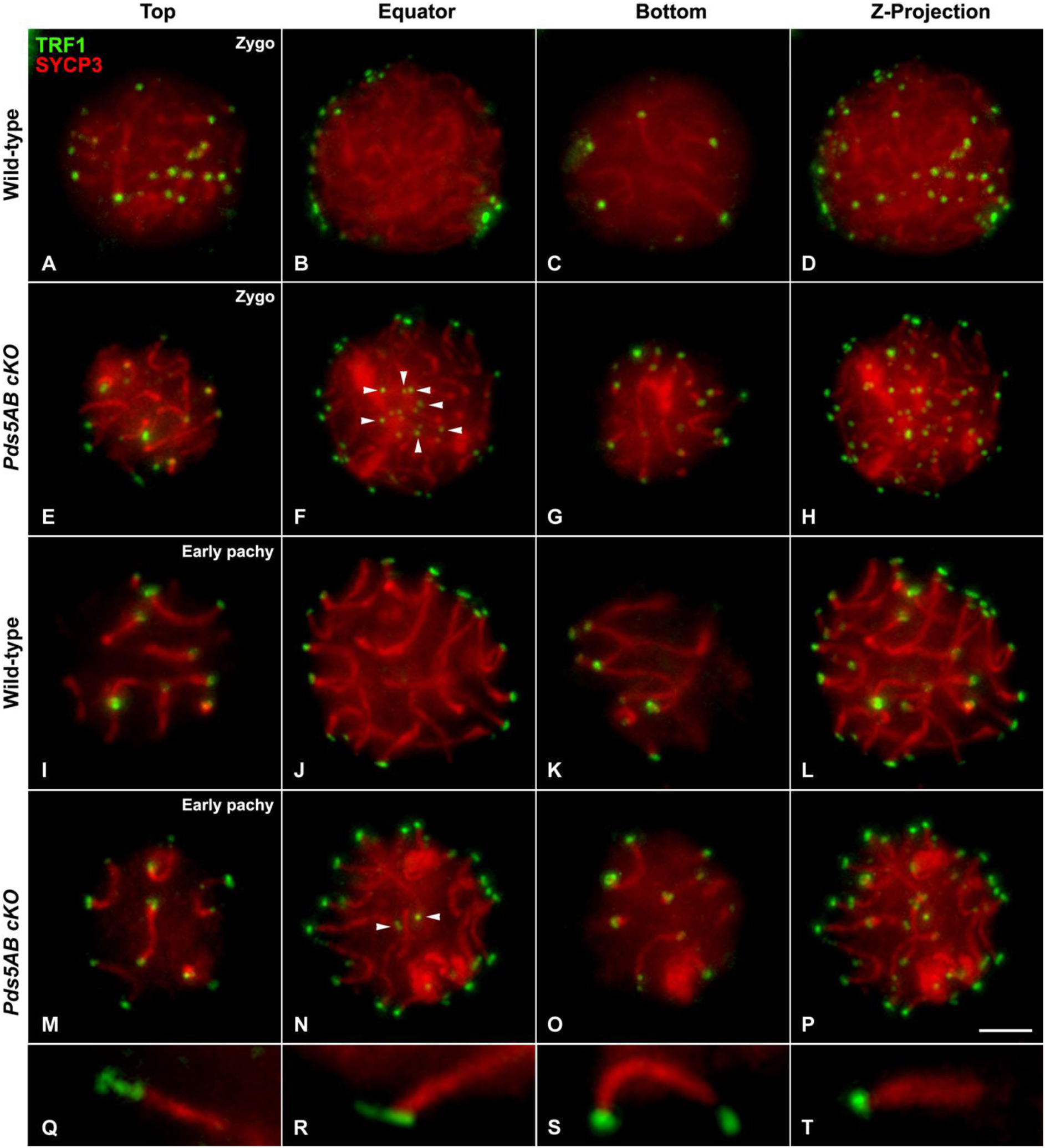
Altered attachment of telomeres to the NE in the absence of PDS5 proteins. **(A-T)** Double immunolabeling of TRF1 (green) and SYCP3 (red) in squashed spermatocytes from wild-type (A-D, I-L) or tamoxifen-treated *Pds5AB* cKO mice (E-H, M-T). The first three columns correspond to z-projections of 15 focal planes throughout the Top, Equator and Bottom regions of the nucleus. The projection of 65 focal planes across the same spermatocyte nucleus is shown at the fourth column (Z-Projection). White arrowheads in F and N indicate the presence of TRF1 signals non-attached to the NE. **(Q-T)** Selected bivalents from *Pds5AB* deficient spermatocytes displaying structural telomere abnormalities as telomere stretches (Q), multiple telomere (R), distant telomere (S) and telomere-less (T). Scale bar: 10 μm.

## Discussion

In the present study we have examined the distribution of the cohesin cofactors PDS5A and PDS5B in mouse spermatocytes and analyzed the consequences of their depletion in order to understand their contributions to mammalian meiosis. As *Pds5A* and *Pds5B* genes are essential for embryonic development, we employed mice carrying conditional knock out alleles for one or both genes in homozygosis and a *Cre* transgene induced by tamoxifen (Carretero et al., 2013). Due to the deleterious effects of eliminating the PDS5 proteins in the whole organism, tamoxifen treatment was limited to two weeks. In this time, depletion of PDS5 proteins was observed in spermatocytes up to mid-pachytene stage. Taking this limitation into account, we have observed that single elimination of either PDS5 protein does not alter progression through prophase I, while elimination of both results in severe phenotypes on AE organization and telomere integrity.

### PDS5A and PDS5B are located at AEs/LEs but their dynamics are different

Immunofluorescence staining with specific antibodies shows that PDS5A and PDS5B are distributed along the AEs/LEs in prophase I mouse spermatocytes, as previously described for the cohesin subunits SMC1α, SMC1β, SMC3, RAD21, RAD21L, REC8 and STAG3 (Ishiguro, 2019). It is likely that the presence of PDS5A and PDS5B at meiotic chromosome axes occurs in the context of the cohesin complex, as reported for PDS5 in different organisms (van Heemst et al., 1999; Zhang et al., 2005; Ding et al., 2006; Jin et al., 2009), and consistent with the finding that PDS5B co-immunoprecipitates with meiosis-specific cohesin subunits REC8 and SMC1β from mouse spermatocyte extracts (Fukuda and Höög, 2010). Likewise, similar distribution patterns have been previously reported in mouse meiosis for the cohesin cofactors WAPL (Zhang et al., 2008; Brieño-Enríquez et al., 2016), and NIPBL and MAU2 (Visnes et al., 2014). The exception is Sororin, which localizes at the central region of the SC (Gómez at al., 2016; Jordan et al., 2017).

We have found intriguing differences between the dynamics of PDS5A and PDS5B during prophase I. While PDS5A appears at AEs/LEs by zygotene and is displaced from them by mid pachytene, PDS5B is visualized at AEs/LEs during all prophase I stages. Moreover, PDS5B is present at telomeres during prophase I. However, since elimination of either PDS5 protein does not result in defects in meiosis progression, at least up to mid-pachytene, the functional significance of the different association/dissociation dynamics of the two PDS5 proteins is unclear. In particular, the lack of telomere defects in *Pds5B* deficient cells suggests that PDS5A can also perform the function of PDS5B required for telomere integrity even if we do not find PDS5A at telomeres either in wild-type or in *Pds5B* deficient spermatocytes. We cannot discard the possibility that the PDS5A antibody used in our study is unable to detect PDS5A at telomeres.

### PDS5 proteins are required for axial element organization

Spermatocytes lacking both PDS5 proteins are able to fulfill complete synapsis, but display severely shortened AEs/LEs in prophase I. A similar shortening of chromosome axes in the absence of PDS5 has been described in other organisms (van Heemst et al., 1999; Wang et al., 2003; Jin et al., 2009; Ding et al., 2016). The extent of SC shortening in *Pds5AB* deficient mouse spermatocytes is comparable, or even greater, to that found in spermatocytes from cohesin knock out mouse models (Biswas et al., 2016; Ward et al., 2016). Consistent with the dependency of AE/LE assembly on previous loading of cohesin complexes along meiotic chromosomes (Eijpe et al., 2000), cohesin axes also appear shortened. Although it is not possible to make a quantitative assessment in our preparations, we confirm that all the cohesin subunits that we tested (SMC1α, SMC1β, SMC3, REC8, RAD21, RAD21L, and STAG3), as well as the cohesin cofactors Sororin and WAPL, were present at the shortened cohesin axes of *Pds5AB* deficient prophase I cells.

Why do cohesin-deficient chromosomes assemble shortened SCs? Descriptions of meiotic chromosome morphology suggest that chromatin loops emanate from a proteinaceous axial core containing cohesin and LE proteins (Zickler and Kleckner, 1999), but the actual mechanism of chromosome assembly remains unknown. In a recent preprint reporting HiC analyses in yeast meiotic chromosomes, Schalbetter et al. (2019 biorxiv) show that the patterns observed are reminiscent of those obtained in interphase mammalian cells after depletion of WAPL or PDS5, suggesting that reduced dissociation dynamics of cohesin may be an important feature of meiotic cohesin complexes (Wutz et al., 2017). The study further proposes that REC8-dependent loop extrusion is reponsible for these patterns and drives chromosome arm compaction while SC formation and synapsis promote additional compaction. Since SC length is restored in *Smc1β^−/−^Sycp3^−/−^*double mutant spermatocytes, cohesin might in fact counteract chromosome axis compaction promoted by the SC (Novak et al., 2008). Whether cohesin complexes mediating cohesion and those driving loop extrusion are the same or the coordination between them remains an open question (Muller et al., 2018; Holzmann 2019). In any case, PDS5 regulates both activities of cohesin, and it is therefore not surprising that its deletion results in defects similar to those reported for meiotic cohesin mutants. We envision that in both scenarios, reduced cohesin or reduced cohesin dynamics in the absence of PDS5, a lower number of longer chromatin loops would be formed and result in shorter AEs/LEs.

### A role for PDS5 proteins in meiotic recombination?

The initiation of meiotic recombination precedes, and mediates, homologous chromosomes synapsis in prophase I. PDS5A and PDS5B are not required for the formation of SPO11-mediated DSBs or their repair. Nonetheless, a reduced number of RAD51 foci was found along the shortened AEs/LEs of *Pds5AB* deficient spermatocytes, as well as of MLH1 foci, which might point to a decrease of recombination rates, since MLH1 is associated to late recombination nodules (Moens et al., 2007). The reduction in RAD51 foci might be a direct consequence of shorter AEs/LEs (Kleckner et al., 2003). Delayed or defective DSB repair, along with SC shortening, has been reported in meiocytes lacking the cohesin subunits REC8 (Bannister et al., 2004; Xu et al., 2005), SMC1β (Revenkova et al., 2004), STAG3 (Fukuda et al., 2014; Hopkins et al., 2014; Llano et al., 2014; Winters et al., 2014) and RAD21L (Herrán et al., 2011). An additional possibility is that RAD51 recruitment or function is impaired in the absence of PDS5 proteins, as *in vitro* assays show that PDS5B interacts physically with RAD51 and stimulates RAD51-mediated DNA strand invasion (Couturier et al., 2016).

### PDS5 proteins secure telomere integrity

In somatic cells, telomere cohesion guarantees proper replication of the telomeres. The absence of PDS5 proteins, or Sororin, or cohesin-SA1, or treatment with low doses of the DNA replication inhibitor aphidicolin, result in altered telomere organization or telomere fragility (Sfeir et al., 2009; Remeseiro et al., 2012; Carretero et al., 2013). Cohesion is established during the premeiotic S-phase, and it is possible that PDS5 deficiency at this time could explain the telomere defects observed in subsequent meiotic prophase I. However, it is important to note that telomere aberrations identical to those reported here have been observed in *Smc1β^−/−^* spermatocytes, while this meiosis-specific cohesin subunit is not expressed in premeiotic cells (Adelfalk et al., 2009). Moreover, an elegant study expressing SMC1α under the control of the *Smc1β* promoter in *Smc1β^−/−^*mice showed restoration of several meiotic phenotypes including SC length while telomere aberrations remained uncorrected (Biswas et al., 2018). It has therefore been suggested that SMC1β may facilitate the arrangement of telomere DNA into the so-called T-loop and/or the association of telomere factors to ensure proper telomere behavior. In this scenario, PDS5 proteins might control SMC1β function(s) at telomeres, despite our observation that they are not necessary for the loading and maintenance of SMC1β or any other cohesin subunit along chromosomes axes during prophase I. At least PDS5B has been shown to co-immunoprecipitate with SMC1β in mouse spermatocytes (Fukuda and Höög, 2010). Further studies will be necessary to elucidate if telomere aberrations found in *Pds5AB* deficient spermatocytes are due to cohesin miss-regulation during premeiotic DNA replication, a failure in SMC1β regulation during meiosis, or both. Nonetheless, our data overall indicate that PDS5 proteins are crucial for the proper arrangement and functionality of telomeres in male mouse meiosis.

## Materials and Methods

### Animals and ethics statement

Mice carrying conditional knock out alleles for Pds5A (Pds5A*^loxfrt^* or *Pds5A* cKO), Pds5B (Pds5B*^lox^*or *Pds5B* cKO) or both (Pds5A*^loxfrt^*; Pds5B*^lox^*or *Pds5AB* cKO) in homozygosis and a Cre-ERT2 transgene, generated previously (Carretero et al., 2013), were used for this study. Animals were kept at the Animal Facility of the Spanish National Cancer Research Centre (CNIO) under specific pathogen-free conditions in accordance with the recommendation of the Federation of European Laboratory Animal Science Associations (FELASA). To induce recombination, 8 week-old male mice were fed with tamoxifen-containing diet for 2 weeks before being sacrificed and their testes surgically removed. All animal procedures were approved by local and regional ethics committees (Institutional Animal Care and Use Committee and Ethics Committee for Research and Animal Welfare, Instituto de Salud Carlos III) and performed according to the European Union guidelines. Genotyping was performed by PCR with the following primers, as indicated in Figure S1A: Afw: 5’-GGACACTTTAGCAGTTACCTCAGC-3’, Arv1: 5’-ACCCTAAGTCCCAATGCACC-3’ and Arv2: 5’-GGCGGAAAGAACCATCTAGC-3’ for Pds5A; Bfw: 5’-GCCCTTCTTTCATTGTTTAC-3’ and Brv: 5’-GGTTTGCAGAGAGTTCTAGC-3’ for Pds5B. Protein levels were also analyzed by immunoblot of total protein extracts prepared by lysing a piece of testis tissue in RIPA buffer. Other knock out mice used in this study were: *Syce3^−/−^* (a gift from R. Benavente; Schramm et al., 2011), *Sycp1^−/−^* (a gift from C. Höög; de Vries et al., 2005), *Rec8^−/−^*(a gift from J.C. Schimenti; Bannister et al., 2004), and *Smc1β^−/−^*(a gift from R. Jessberger; Revenkova et al., 2004).

### Squashing and spreading of spermatocytes

For immunofluorescence analyses, testes were detunicated and seminiferous tubules processed according to previously described protocols for obtaining squashing (Page et al., 1998; Parra et al., 2002) or spreading (Peters et al., 1997) preparations of spermatocytes.

### Immunofluorescence microscopy

After spermatocytes processing, the resulting preparations were rinsed three times for 5 min in PBS, and incubated overnight at 4°C with the corresponding combinations of primary antibodies diluted in PBS. A rabbit polyclonal antibody for mouse PDS5A (C-107) was generated by immunizig rabbits with a recombinant fragment (aa. 1188-1332) expressed in *Escherichia coli*. Two rabbit polyclonal antibodies were prepared against mouse PDS5B: for C-100 a synthetic peptide (aa. 2-24, HSKTRTNDGKITYPPGVKEISD) was used as immunogen; for C-102 a recombinant fragment (aa. 1178-1300) expressed in *E. coli* was used. Both antibodies against PDS5B yielded identical results in immunofluorescence stainings of testes. An antibody against human PDS5B prepared by immunizing rabbits with a synthetic peptide (CEEKLGMDDLTKLVQEQKPKGSQRS, aa. 1226-1249) was affinity purified and used for western blot. All primary antibodies and dilutions used are presented in Table S2. Following three washes in PBS, the slides were incubated for 45 min at room temperature with secondary antibodies at a 1:100 dilution in PBS. The appropriated combinations of the following secondary antibodies were employed for simultaneous double- or triple-immunolabeling: Alexa 350, Alexa 488 and Alexa 594-conjugated donkey anti-rabbit IgG (Molecular Probes); Alexa 350, Alexa 594 and Alexa 594-conjugated donkey anti-mouse IgG (Molecular Probes); Alexa 488-conjugated donkey anti-goat IgG (Molecular Probes); Alexa 488-conjugated goat anti-guinea pig IgG (Molecular Probes); Alexa 488-conjugated donkey anti-human IgG (Molecular Probes). Subsequently, slides were rinsed in PBS, and in double-immunolabeling experiments counterstained for 3 min with 5 μg/ml DAPI (4’, 6-diamidino-2-phenylindole). After a final rinse in PBS, the slides were mounted in Vectashield (Vector Laboratories) and sealed with nail varnish.

### Immunofluorescence-FISH

A combination of SYCP3 immunostaining, our double immunolabeling of SYCP3 and RAP1, and telomere FISH were carried out in spread spermatocytes as previously reported (Viera et al., 2003). For this purpose, a FITC-labeled (C3TA2)3 peptide nucleic acid (PNA) probe (Applied Biosystems) that specifically recognizes telomeric DNA was employed.

### Histology

For histological sections, testes were divided in several pieces, which were fixed by immersion in Bouin’s solution for 24 hours. After standard washes and dehydration, Paraplast-embedded tissue blocks were cut in 3 μm thick sections in a Reichert microtome. Finally, sections were stained with conventional Mallory’s trichrome stain.

### Image acquisition

Observations were performed using an Olympus BX61 microscope equipped with a motorized Z axis and epifluorescence optics. Images were captured with an Olympus DP71 digital camera controlled by the Cell^P^ software (Olympus). For squashed spermatocytes, image stacks comprising 65 focal planes across the entire spermatocyte nuclei were captured. The complete image stacks were used to analyze the relative distribution of the proteins at a given entire nucleus and for generating animated 3D reconstructions of nuclei using the public domain software ImageJ (National Institutes of Health, USA; http://rsb.info.nih.gov/ij). For a better presentation of the results in the figures, stacks were subsequently processed for obtaining complete Z-projections (65 focal planes) or partial Z-projections (15 focal planes along a certain nuclear region) using the ImageJ software. Final images were processed with Adobe Photoshop 7.0 software.

### Quantification and statistical analyses

To measure SC length, each autosomal SC was scored in 33 early pachytene cells at random, in preparations from three different individuals of each genotype, using the “measurement” function of the ImageJ software. The SC length of the sex chromosomes was disregarded since these chromosomes do not achieve complete synapsis and might introduce errors. Since no statistical differences were found between individuals within a given genotype by a Kruskal-Wallis test, the data were grouped and analyzed by a T-test function (confidence level: 95%) using GraphPad Prism 6.0 software.

In order to determine the percentage of spermatocytes in a given prophase I sub-stage, we analyzed and classified 1,000 spermatocytes at random in three individuals for each genotype. For this purpose, spermatocytes were double immunolabeled for SYCP3 and SYCP1 to determine the progression of synapsis, and thus the meiotic staging, of each prophase I spermatocyte.

The quantification of RAD51 foci was performed in 15 zygotene nuclei, randomly selected, in preparations from three different individuals of each genotype using the “multi-point” function of the ImageJ software. It is worth noting that only RAD51 foci located over the LEs were scored. Since no statistical differences were found between individuals within a given genotype by a Kruskal-Wallis test, the data were grouped and analyzed by a T-test function (confidence level: 95%) using GraphPad Prism 6.0 software. A similar approach was employed to quantify MLH1 foci in 100 early pachytene spermatocytes displaying shortened SCs and in 91 mid/late pachytene spermatocytes with regular SCs.

Telomere morphology was evaluated after the immunolabelling of the telomere associated proteins TRF1, RAP1 and SUN1 and telomeric DNA FISH. Anomalous telomere signals were classified as (i) telomere stretches: stretched signals emerging from the AE/LE ends; (ii) multiple telomere: multiple signals at a given AE/LE end; (iii) distant telomeres: a single signal separated from an AEs/LEs end; and (iv) telomere-less: AEs/LEs ends without a detectable signal. For TRF1 we scored 1646 telomeres in zygotene spermatocytes and 738 in pachytene ones. For RAP1 we evaluated 1376 and 328 telomeres in zygotene and early pachytene spermatocytes, respectively. A total amount of 715 zygotene and 410 early pachytene telomeres were analysed to evaluate SUN1 morphology. Telomere FISH signals were examined in 1198 zygotene telomeres and in 574 pachytene ones.

## Acknowledgements

We express our sincere thanks to Ricardo Benavente, Christer Höög, John Schimenti and Rolf Jessberger for providing SYCE3, SYCP1, REC8 and SMC1β knock out male mice, respectively; and to Jibak Lee, Mary Ann Handel, Attila Tóth and Manfred Alsheimer for providing antibodies. We also thank Lorena Barreras for expert histological assistance and Miriam Rodríguez for mouse genotyping. This work was supported by Ministerio de Economía y Competitividad (Spain) and FEDER funds through grants BFU2014-53681-P (to JAS), BFU2016-79841-R (to AL), BFU2017-89408-R, FPI “Severo Ochoa” fellowship to M.R.-T. and Centro de Excelencia “Severo Ochoa” to CNIO.

## Author contributions

AV, AL and JAS conceived the study; AV, IB, MR-T, RG, AG and JAS performed all experiments; JLB developed PDS5 antibodies C-100, C-102 and C-107; AV, AL and JAS analyzed results and wrote the paper with some contributions from the other co-authors.

## Supplementary figure legends

**Figure S1.**
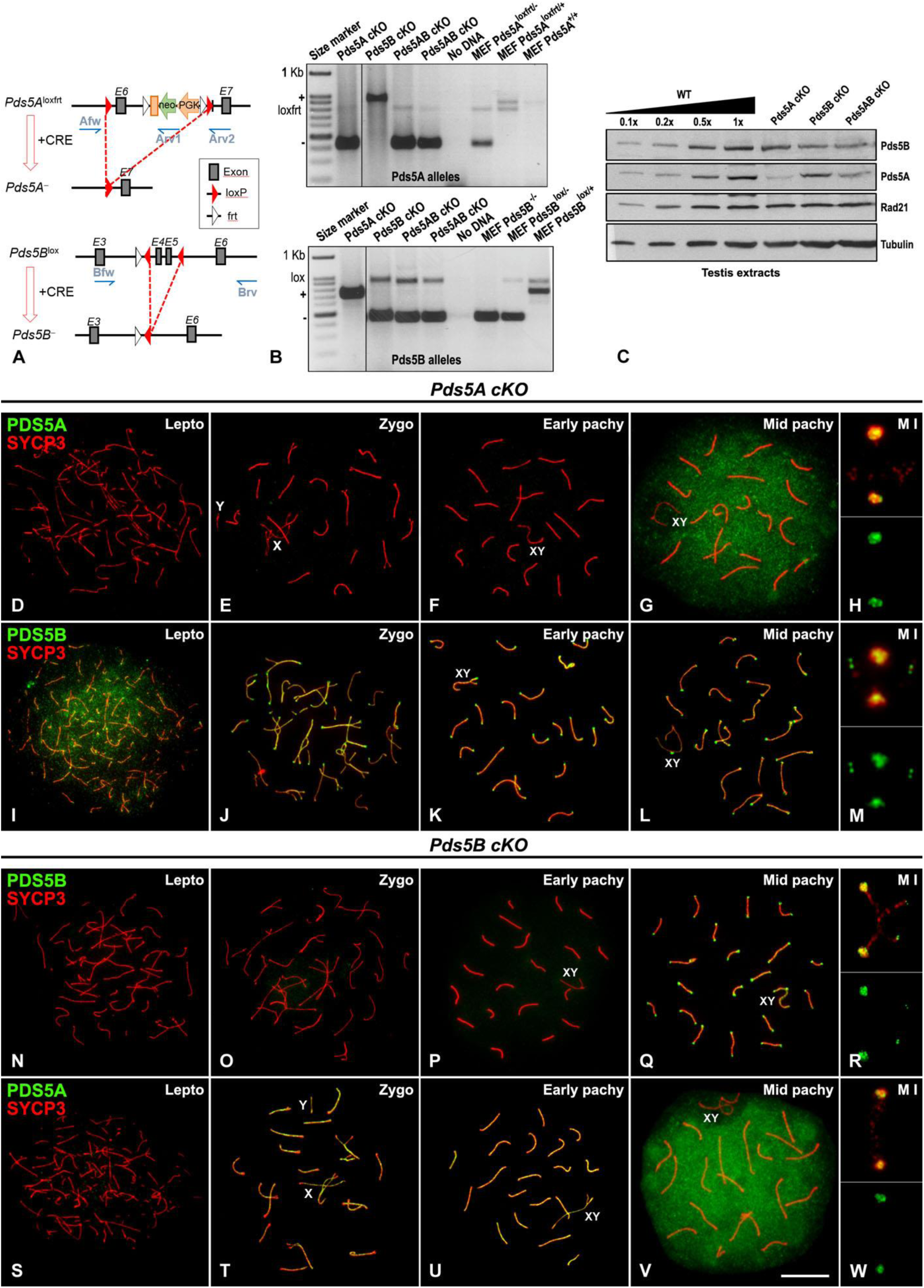
PDS5A and PDS5B elimination in spermatocytes. **(A)** Schematic representation of the conditional alleles (*loxfrt* or *lox*) and the null allele (−) obtained upon Cre-mediated recombination for *Pds5A* (top) and *Pds5B* (bottom). The position of the primers used for genotyping is indicated (blue arrows). **(B)** Example of genotyping PCRs for *Pds5A* (top) and *Pds5B* (bottom) alleles using DNA obtained from testis of the indicated mice as well as DNA from mouse embryo fibroblasts as control. cKO mice carry the conditional allele(s) in homozygosis and a Cre-ERT2 transgene and have been treated with tamoxifen to promote the translocations of the Cre recombinase to the nucleus. In most cKO mice, excision of the targeted exon is not complete and a weak band corresponding to the conditional allele can be still be detected in addition to the stronger band of the null allele. The sizes of the different PCR products are: *Pds5A* wild-type (+) 872 bp, loxfrt 778 bp and null (−) 414 bp; for *Pds5B* wild-type (+) 706 bp, lox 859 bp and null (−) 415 bp. **(C)** Immunoblot analysis of total extracts prepared from testes of mice of the indicated genotypes. Decreasing amounts of extract from testes obtained from wild-type mice (WT) were loaded for comparison. Overall, elimination of *Pds5B* is less efficient than elimination of *Pds5A* both in the single and in the double cKO mice. **(D-M)** Double immunolabeling of PDS5A (green, D-H) or PDS5B (green, I-M) and SYCP3 (red) in spread spermatocytes from tamoxifen-treated *Pds5A* cKO in leptotene (D; Lepto), zygotene (E; Zygo), early pachytene (F; Early pachy), mid pachytene (G; Mid pachy) and in a metaphase I bivalent (H; M I). **(N-W)** As above, but in spermatocytes from tamoxifen-treated *Pds5B* cKO mice. In both cases, by mid-pachytene PDS5 proteins are not depleted. The position of sex bivalents (XY) is indicated. Scale bar: 10 μm.

**Figure S2.**
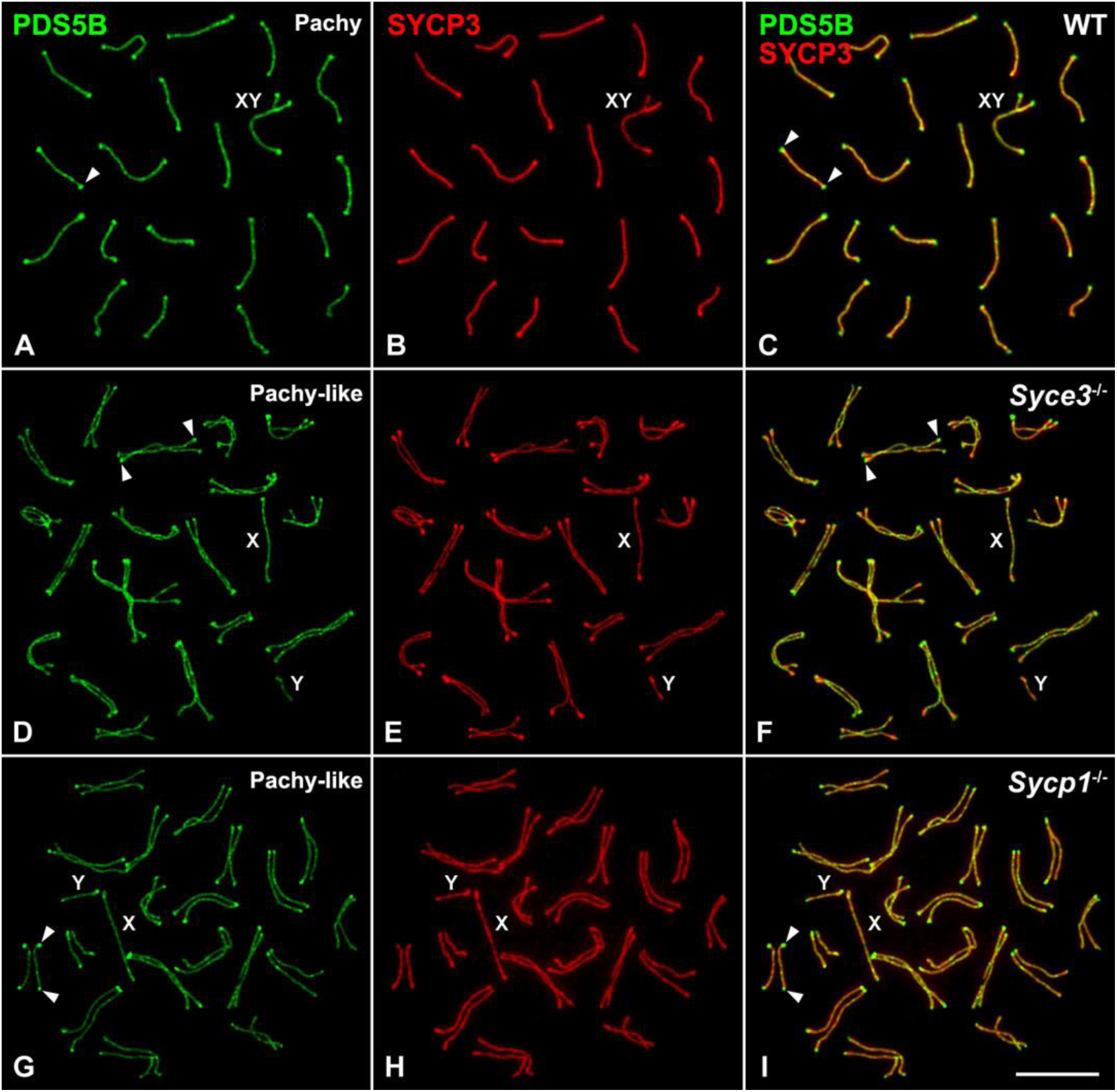
PDS5B is enriched at the ends of aligned but non-synapsed AEs/LEs. **(A-I)** Double immunolabeling of PDS5B (green) and SYCP3 (red) in a spread wild-type (WT) pachytene spermatocyte (A-C; Pachy), and *Syce3^−/−^* (D-F) and *Sycp1^−/−^* (G-I) pachytene-like (Pachy-like) spermatocytes. Arrowheads indicate the accumulation of PDS5B at the ends of SCs and aligned LEs. The sex bivalent (XY) and sex chromosomes (X, Y) are also indicated. Scale bar: 10 μm.

**Figure S3.**
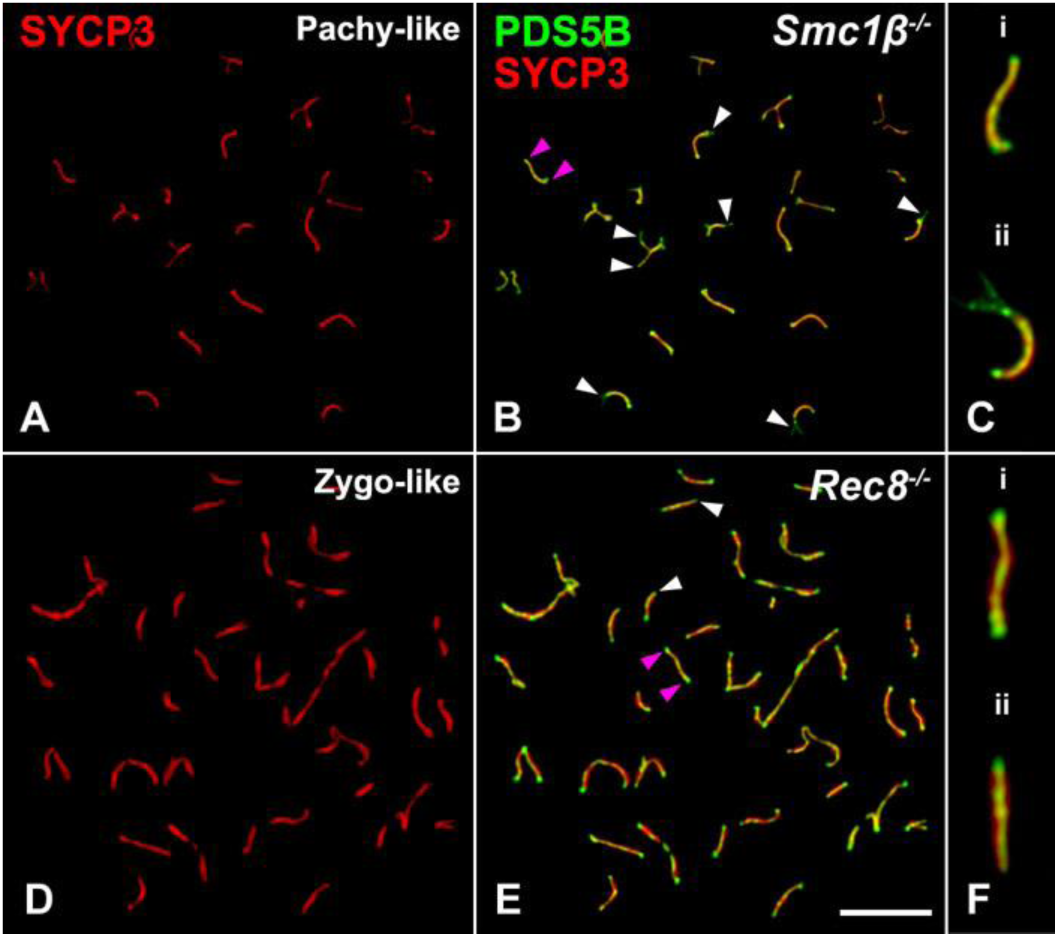
PDS5B is present at altered telomeres in Smc1β−/− and Rec8−/− spermatocytes. **(A-F)** Double immunolabeling of PDS5B (green) and SYCP3 (red) in spread pachytene-like *Smc1β^−/−^* (A-C) and zygotene-like *Rec8^−/−^*(D-F) spermatocytes. White arrowheads in B and E indicate telomeres with absence or altered accumulation of PDS5B. Pink arrowheads in B and E indicate telomeres with a regular PDS5B accumulation. Selected pachytene-like bivalents displaying regular accumulation of PDS5B at telomeres (i) or structural telomere abnormalities (ii). Scale bar: 10 μm.

**Figure S4.**
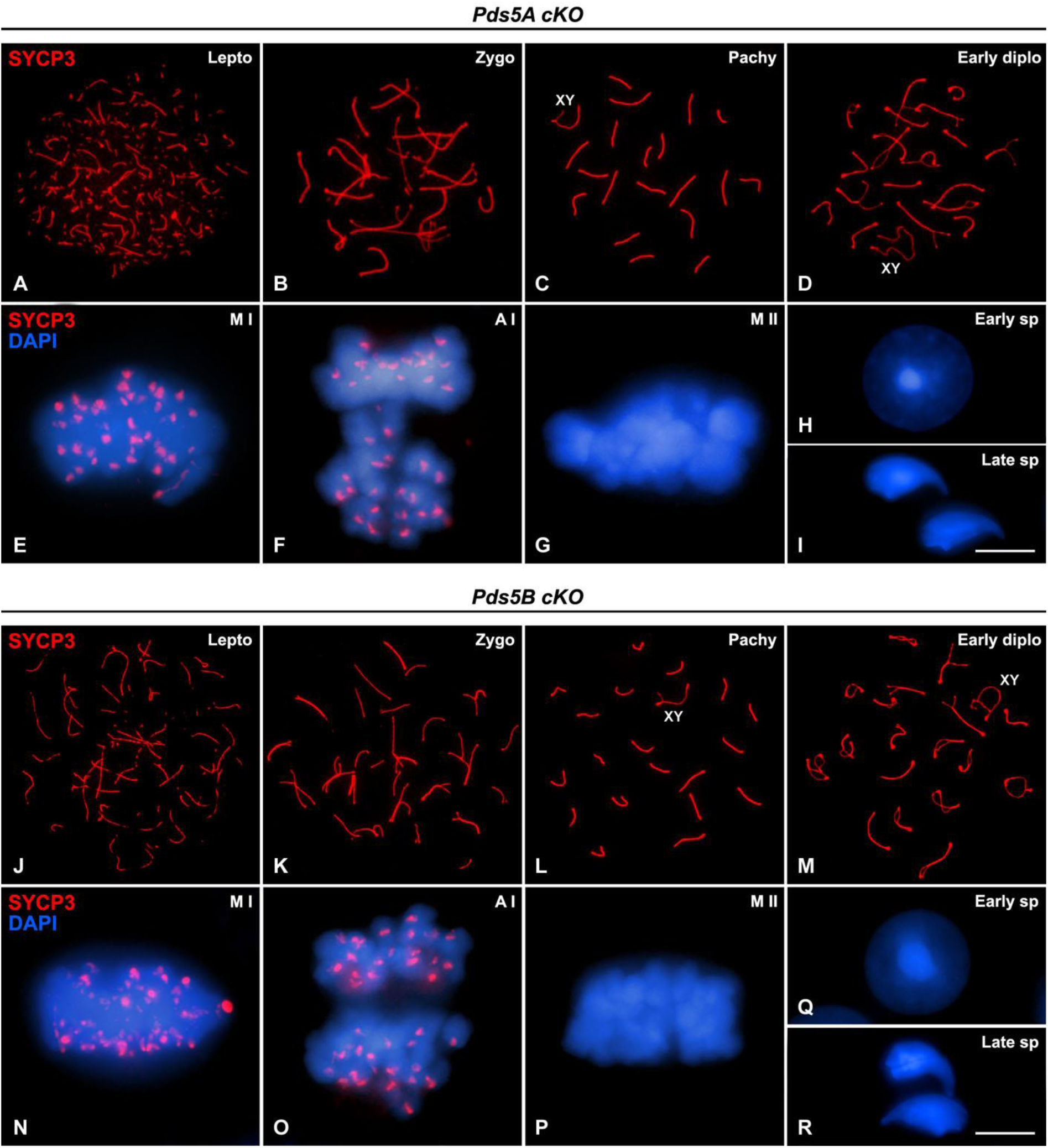
Spermatogenesis is not altered in *Pds5A* cKO or *Pds5B* cKO mice. **(A-G)** Immunolabeling of SYCP3 (red) in spread (A-D) and squashed (E-G) *Pds5A* cKO spermatocytes in the stages indicated. DAPI staining of chromatin (blue) is shown in E-G. **(H and I)** DAPI stained early (H; Early sp) and late spermatid nuclei (I; Late sp) from *Pds5A* cKO mice treated with tamoxifen. **(J-P)** Immunolabeling of SYCP3 (red) in spread (J-M) and squashed (N-P) *Pds5B* cKO spermatocytes in the stages indicated. DAPI staining of chromatin (blue) is shown in N-P. **(Q and R)** DAPI stained early (Q; Early sp) and late spermatid nuclei (R; Late sp) from *Pds5B* cKO mice treated with tamoxifen. The position of sex bivalents (XY) is indicated. Scale bar: 10 μm.

**Figure S5.**
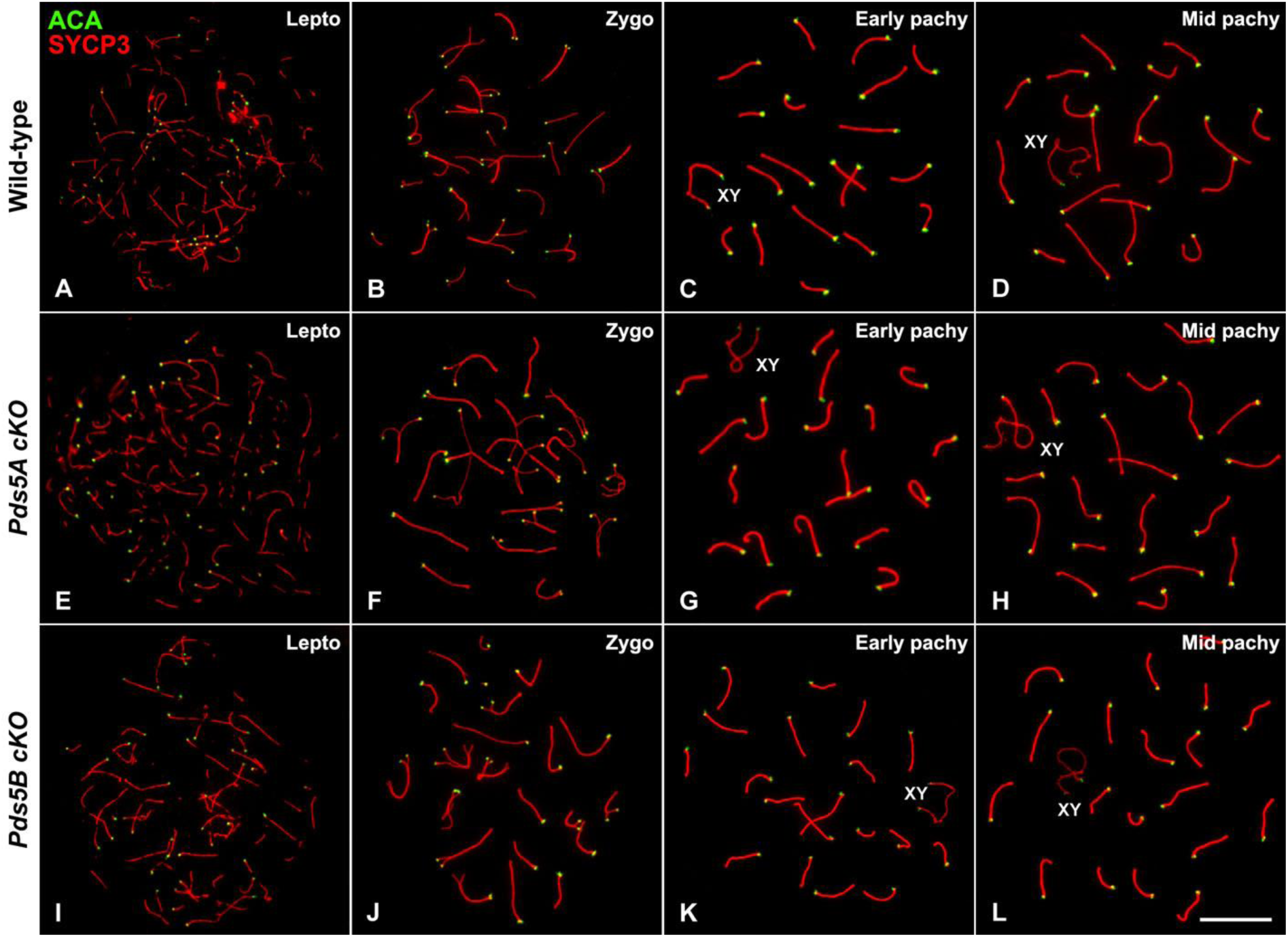
A single PDS5 protein supports proper centromere organization in meiotic chromosomes before mid-pachytene. **(A-L)** Double immunolabeling of kinetochores (ACA, green) and SYCP3 (red) in spread spermatocytes from wild-type (A-D), tamoxifen-treated *Pds5A* cKO (E-H) or *Pds5B* cKO (I-L) mice in the indicated stages. The position of sex bivalents (XY) is indicated. Scale bar: 10 μm.

**Figure S6.**
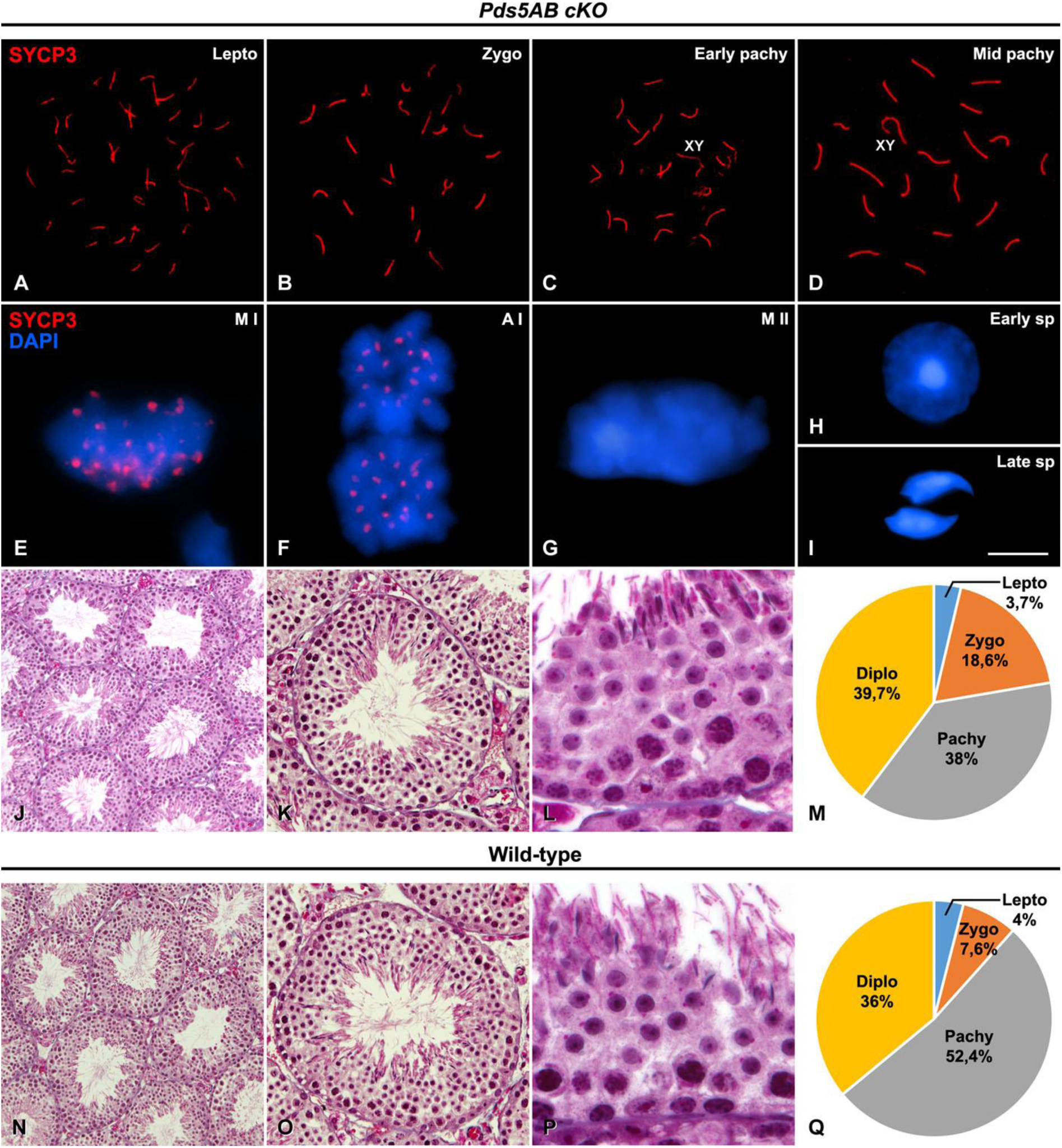
Spermatogenesis progression in *Pds5AB* cKO mice. **(A-G)** Immunolabeling of SYCP3 (red) in spread (A-D) and squashed (E-G) spermatocytes from tamoxifen-treated double *Pds5AB* cKO mice in the stages indicated. DAPI staining (blue) is shown in E-G. **(H and I)** DAPI stained early (H; Early sp) and late spermatid nuclei (I; Late sp). The position of sex bivalents (XY) is indicated. Scale bar: 10 μm. **(J-L and N-P)** Histological sections of testes from the same mice (J-L) as well as wild-type (N-P). **(M and Q)** Pie graph showing the distribution of prophase I sub-stages after analyzing 1,000 spermatocytes from three individuals of each genotype double immunolabeled for SYCP3 and SYCP1.

**Figure S7.**
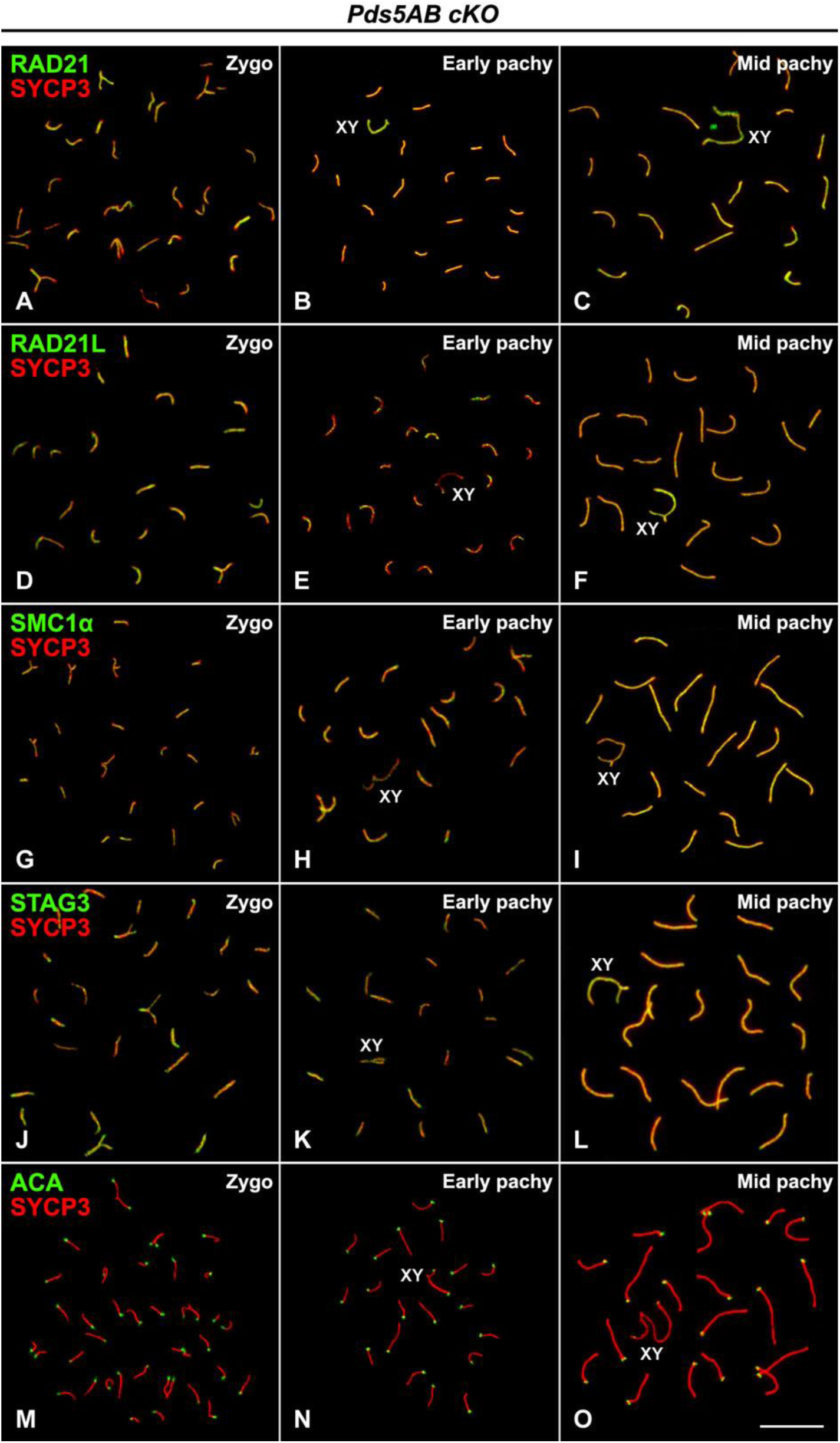
Distribution of cohesin subunits in the absence of PDS5A and PDS5B. **(A-L)** Double immunolabeling of SYCP3 (red) with the cohesin subunits RAD21 (green in A-C), RAD21L (green in D-F), SMC1α (green in G-I) and STAG3 (green in J-L) in spermatocytes from tamoxifen-treated *Pds5AB* cKO mice in the indicated stages. **(M-O)** Double immunolabeling of kinetochores (ACA, green) and SYCP3 (red) in the same spermatocytes. The position of sex bivalents (XY) is indicated. Scale bar: 10 μm.

**Figure S8.**
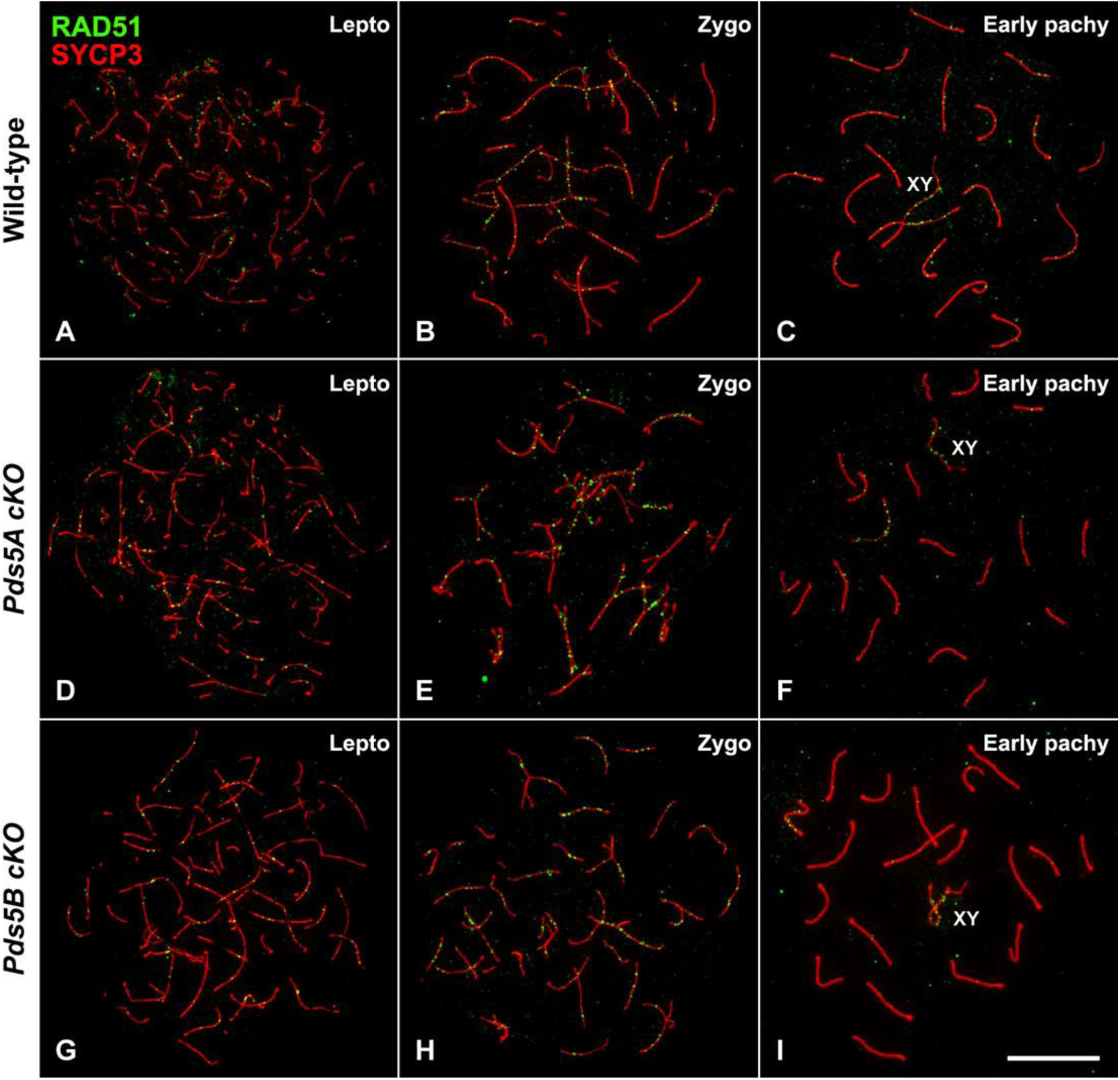
A single PDS5 protein is sufficient for proper RAD51 localization in early prophase I spermatocytes. **(A-L)** Double immunolabeling of RAD51 (green) and SYCP3 (red) in spread leptotene (A, D and G; Lepto), zygotene (B, E and H; Zygo) and early pachytene (C, F and I; Early pachy) spermatocytes from wild-type (A-C) and tamoxifen-treated *Pds5A* cKO (D-F) and *Pds5B* cKO individuals (G-I). The position of sex bivalents (XY) is indicated. Scale bar: 10 μm.

**Figure S9.**
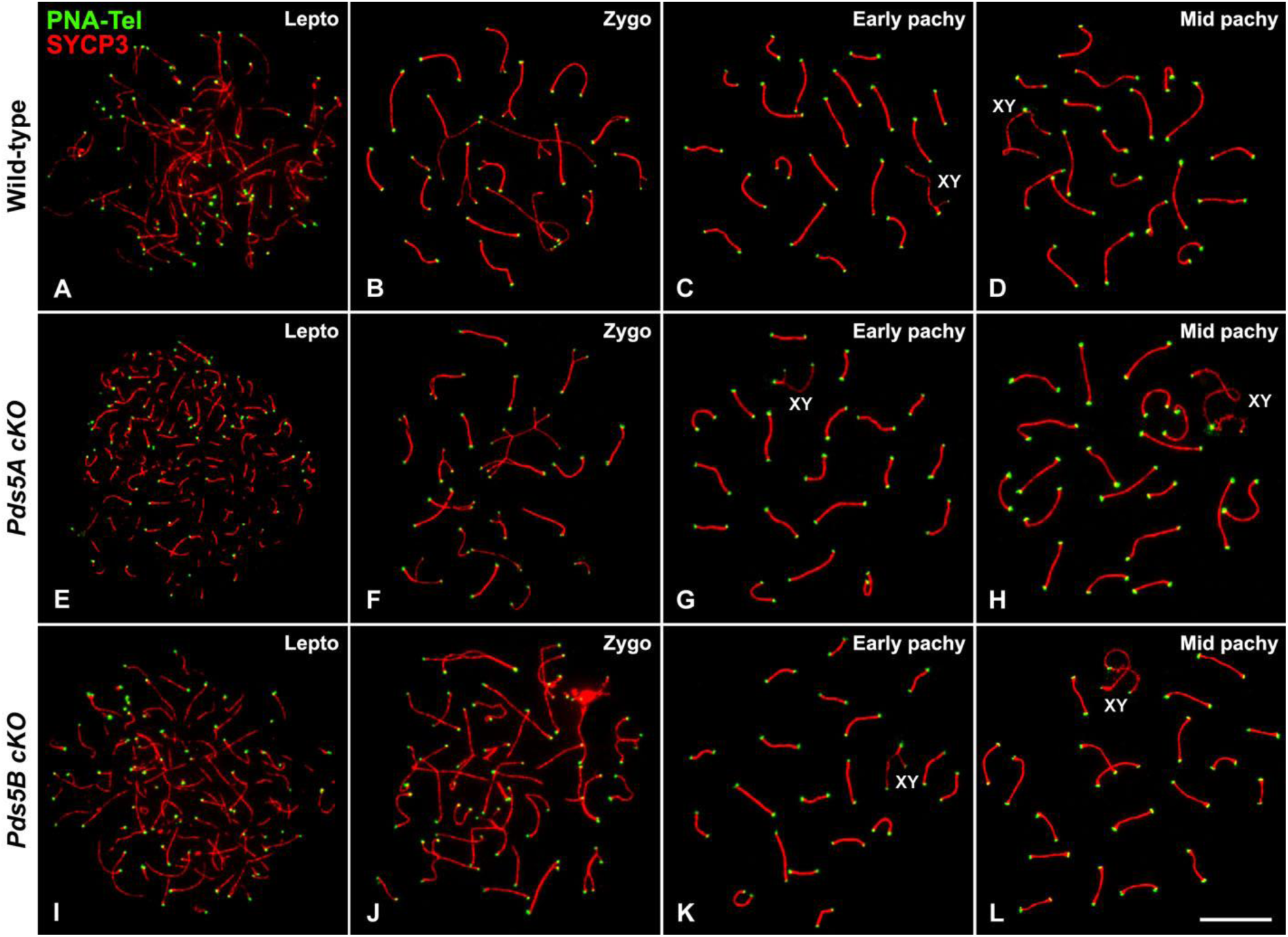
Telomeric DNA organization is not altered in the absence of a single PDS5 protein. **(A-L)** Immunolabeling of SYCP3 (red) and telomere FISH (PNA-Tel, green) in spread spermatocytes from wild-type (A-D), and tamoxifen-treated *Pds5A* cKO (E-H) and *Pds5B* cKO (I-L) mice in the indicated stages. The position of sex bivalents (XY) is indicated. Scale bar: 10 μm.

**Figure S10.**
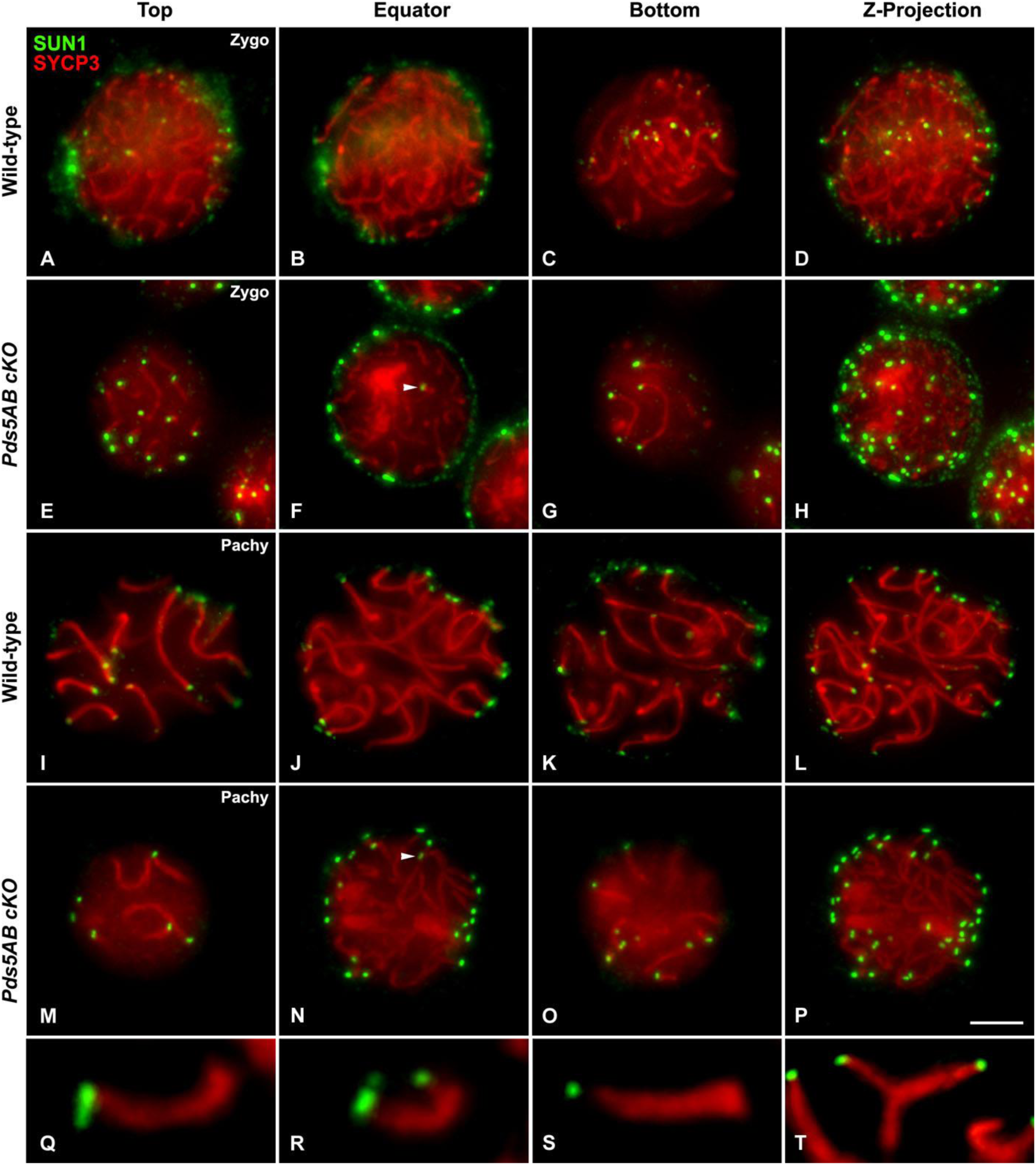
Defects in SUN1 distribution and telomere attachment to the NE in the absence of PDS5 proteins. **(A-T)** Double immunolabeling of SUN1 (green) and SYCP3 (red) in squashed zygotene (A-H; Zygo) and early pachytene (I-P; Early pachy) spermatocytes in wild-type (A-D, I-L) and tamoxifen-treated *Pds5AB* cKO mice (E-H, M-T). The first three columns correspond to z-projections of 15 focal planes throughout the Top, Equator and Bottom regions of the nucleus. The projection of 65 focal planes across the same spermatocyte nucleus is shown at the fourth columns (Z-Projection). White arrowheads in F and N indicate the presence of SUN1 signals telomeres non-attached to the NE. **(Q-T)** Selected zygotene/pachytene bivalents from Pds5AB deficient spermatocytes displaying structural telomere abnormalities as telomere stretches (Q), multiple telomere (R), distant telomere (S) and telomere-less (T). Scale bar: 10 μm.

## Supplementary table legends

**Table S1.**
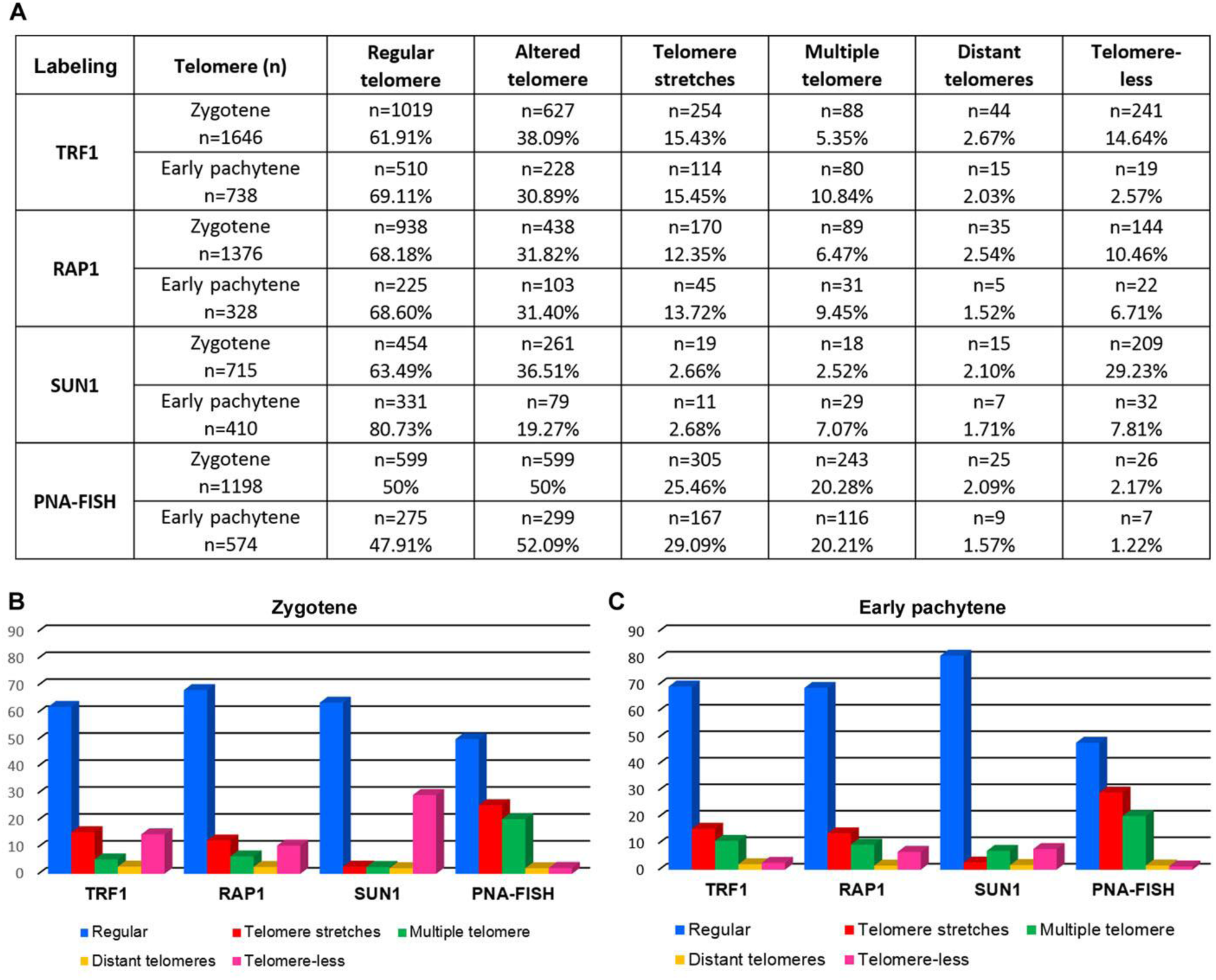
Telomere defects in the absence of PDS5 proteins. **(A)** Quantification of telomere morphology in zygotene and early pachytene spermatocytes from tamoxifen treated *Pds5AB* cKO mice after the immunolabeling of TRF1, RAP1 and SUN1, or FISH for telomeric repeats (PNA-FISH). Altered telomeres are classified as telomere stretches, multiple telomere, distant telomeres or telomere-less (see text for further information). **(B and C)** Bar graph representation of the data in (A).

| Labeling | Telomere (n) | Regular telomere | Altered telomere | Telomere stretches | Multiple telomere | Distant telomeres | Telomere-less |
| --- | --- | --- | --- | --- | --- | --- | --- |
| TRF1 | Zygotene<br>n=1646 | n=1019<br>61.91% | n=627<br>38.09% | n=254<br>15.43% | n=88<br>5.35% | n=44<br>2.67% | n=241<br>14.64% |
|  | Early pachytene<br>n=738 | n=510<br>69.11% | n=228<br>30.89% | n=114<br>15.45% | n=80<br>10.84% | n=15<br>2.03% | n=19<br>2.57% |
| RAP1 | Zygotene<br>n=1376 | n=938<br>68.18% | n=438<br>31.82% | n=170<br>12.35% | n=89<br>6.47% | n=35<br>2.54% | n=144<br>10.46% |
|  | Early pachytene<br>n=328 | n=225<br>68.60% | n=103<br>31.40% | n=45<br>13.72% | n=31<br>9.45% | n=5<br>1.52% | n=22<br>6.71% |
| SUN1 | Zygotene<br>n=715 | n=454<br>63.49% | n=261<br>36.51% | n=19<br>2.66% | n=18<br>2.52% | n=15<br>2.10% | n=209<br>29.23% |
|  | Early pachytene<br>n=410 | n=331<br>80.73% | n=79<br>19.27% | n=11<br>2.68% | n=29<br>7.07% | n=7<br>1.71% | n=32<br>7.81% |
| PNA-FISH | Zygotene<br>n=1198 | n=599<br>50% | n=599<br>50% | n=305<br>25.46% | n=243<br>20.28% | n=25<br>2.09% | n=26<br>2.17% |
|  | Early pachytene<br>n=574 | n=275<br>47.91% | n=299<br>52.09% | n=167<br>29.09% | n=116<br>20.21% | n=9<br>1.57% | n=7<br>1.22% |

**Table S2.** Primary antibodies used in this study.

| **Antigen** | **Host** | **Source or reference** | **Cat. Number** | **IF dilution** | **WB dilution** |
| --- | --- | --- | --- | --- | --- |
| mPDS5A | Rabbit | C-107<br>Custom-made<br>aa. 1188-1332 | -- | 1:10 | -- |
| mPDS5A | Rabbit | Carretero et al., 2013 | -- | -- | 2 µg/ml |
| mPDS5B | Rabbit | C-100<br>Custom-made<br>aa. 2-24 | -- | 1:10 | -- |
| hPDS5B | Rabbit | C-102<br>Custom-made<br>aa. 1178-1300 | -- | 1:10 | -- |
| hPDS5B | Rabbit | Custom-made<br>aa. 1226-1249 | -- | -- | 4 µg/ml |
| mSYCP3 | Mouse | Santa Cruz | sc-74569 | 1:100 | -- |
| mSYCP1 | Goat | Santa Cruz | sc-20837 | 1:20 | -- |
| Kinetochores (ACA) | Human | Antibodies Inc. | 15-234 | 1:20 | -- |
| hSMC1α | Rabbit | K988<br>Krasilova et al., 2005 | -- | 1:5 | -- |
| hSMC1β | Rabbit | K974<br>Prieto et al., 2004 | -- | 1:5 | -- |
| hSMC3 | Rabbit | Abcam | ab-9263 | 1:10 | -- |
| mRAD21 | Rabbit | Carretero et al., 2013 | -- | -- | 2 µg/ml |
| hRAD21 | Rabbit | K854<br>Prieto et al., 2002 | -- | 1:5 | -- |
| mRAD21L | Rabbit | Herrán et al., 2011 | -- | 1:10 | -- |
| mREC8 | Rabbit | Lee et al., 2003 | -- | 1:10 | -- |
| hSTAG3 | Rabbit | K403<br>Prieto et al., 2001 | -- | 1:10 | -- |
| mSororin | Rabbit | C-106<br>Gómez et al., 2016 | -- | 1:10 | -- |
| hWAPL | Rabbit | Proteintech | 16370-1-AP | 1:10 | -- |
| mTRF1 | Rabbit | Alpha Diagnostic Int. | TRF12-S | 1:50 | -- |
| RAP1 | Rabbit | Imgenex | IMG-289 | 1:50 | -- |
| mSUN1 | Guinea pig | Adelfalk et al., 2009 | -- | 1:30 | -- |
| H1t | Guinea pig | Inselman et al., 2003 | -- | 1:100 | -- |
| γ-H2A.X | Mouse | Millipore | 05-636 | 1:1000 | -- |
| mHORMAD2 | Rabbit | Wojtasz et al., 2009 | -- | 1:50 | -- |
| hRAD51 | Rabbit | Merck | Ab-1 PC130 | 1:5 | -- |
| hMLH1 | Mouse | Pharmingen | 511091 | 1:5 | -- |
| Tubulin | Mouse | Clone DM1A, Sigma | T6199 | -- | 1:1000 |

## Supplementary movies legends

**Movie S1.** 3D reconstructions of a complete nucleus (left) and its equatorial region (right) of a squashed pachytene spermatocyte from a tamoxifen-treated *Pds5AB* cKO mouse immunolabeled for SYCP3 (red) and TRF1 (green). The white arrow indicates the position of a telomere non-attached to the NE.

**Movie S2.** 3D reconstruction of the complete nucleus shown in Figure 10Q, of a squashed pachytene spermatocyte from a tamoxifen-treated *Pds5AB* cKO mouse immunolabeled for SYCP3 (red) and TRF1 (green). The white arrow indicates the position of telomere stretches.

**Movie S3.** 3D reconstruction of the complete nucleus shown in Figure 10R, of a squashed pachytene spermatocyte from a tamoxifen-treated *Pds5AB* cKO mouse immunolabeled for SYCP3 (red) and TRF1 (green). The white arrow indicates the position of a multiple telomere.

**Movie S4.** 3D reconstruction of the complete nucleus shown in Figure 10S, of a squashed pachytene spermatocyte from a tamoxifen-treated *Pds5AB* cKO mouse immunolabeled for SYCP3 (red) and TRF1 (green). The white arrow indicates the position of a distant telomere.

**Movie S5.** 3D reconstructions of a complete nucleus (left) and its equatorial region (right) of a squashed pachytene spermatocyte from a tamoxifen-treated *Pds5AB* cKO mouse immunolabeled for SYCP3 (red) and TRF1 (green) shown in Figure 10T. The white arrow indicates the position of telomere-less SC end.

**Movie S6.** 3D reconstructions of a complete nucleus (left) and its equatorial region (right) of a squashed pachytene spermatocyte from a tamoxifen-treated *Pds5AB* cKO mouse immunolabeled for SYCP3 (red) and SUN1 (green) shown in Figure S10E-H. The white arrow indicates the position of a telomere non-attached to the NE and labeled for SUN1.

**Movie S7.** 3D reconstructions of a complete nucleus (left) and its equatorial region (right) of a squashed pachytene spermatocyte from a tamoxifen-treated *Pds5AB* cKO mouse immunolabeled for SYCP3 (red) and SUN1 (green) shown in Figure S10M-P. The white arrow indicates the position of a telomere non-attached to the NE and labeled for SUN1.

**Movie S8.** 3D reconstruction of the equatorial region of a squashed pachytene spermatocyte shown in Figure S10T immunolabeled for SYCP3 (red) and SUN1 (green). The white arrow indicates the position of a SC end with no SUN1 labeling that is not attached to the NE.

## References

Adelfalk C, Janschek J, Revenkova E, Blei C, Liebe B, Göb E, Alsheimer M, Benavente R, de Boer E, Novak I et al (2009) Cohesin SMC1β protects telomeres in meiocytes. J Cell Biol 187: 185–199

Bannister LA, Reinholdt LG, Munroe RJ, Schimenti JC (2004) Positional cloning and characterization of mouse mei8, a disrupted allelle of the meiotic cohesin Rec8. Genesis 40: 184–194

Biswas U, Wetzker C, Lange J, Christodoulou EG, Seifert M, Beyer A, Jessberger R (2013) Meiotic cohesin SMC1β provides prophase I centromeric cohesion and is required for multiple synapsis-associated functions. PLoS Genet 9: e1003985

Biswas U, Hempel K, Llano E, Pendás A, Jessberger R (2016) Distinct roles of meiosis-specific cohesin complexes in mammalian spermatogenesis. PLoS Genet 12: e1006389

Biswas U, Stevense M, Jessberger R (2018) SMC1α substitutes for many meiotic functions of SMC1β but cannot protect telomeres from damage. Curr Biol 28: 249–261

Bolcun-Filas E, Handel MA (2018) Meiosis: the chromosomal foundation of reproduction. Biol Reprod 99: 112–126

Brieño-Enríquez MA, Moak SL, Toledo M, Filter JJ, Gray S, Barbero JL, Cohen PE, Holloway JK (2016) Cohesin removal along the chromosome arms during the first meiotic division depends on a NEK1-PP1γ-WAPL axis in the mouse. Cell Rep 17: 977–986

Carretero M, Ruíz-Torres M, Rodríguez-Corsino M, Barthelemy I, Losada A (2013) Pds5B is required for cohesion establishment and Aurora B accumulation at centromeres. EMBO J 32: 2938–2949

Chicheportiche A, Bernardino-Sgherri J, de Massy B, Dutrillaux B (2007) Characterization of Spo11-dependent and independent phospho-H2AX foci during meiotic prophase I in the male mouse. J Cell Sci 120: 1733–1742

Couturier AM, Fleury H, Patenaude AM, Bentley VL, Rodrigue A, Coulombe Y, Niraj J, Pauty J, Berman JN, Dellaire G et al (2016) Roles for APRIN (PDS5B) in homologous recombination and in ovarian cancer prediction. Nucleic Acids Res 44: 10879–10897

de Vries FA, de Boer E, van den Bosch M, Baarends WM, Ooms M, Yuan L, Liu JG, van Zeeland AA, Heyting C, Pastink A (2005) Mouse Sycp1 functions in synaptonemal complex assembly, meiotic recombination, and XY body formation. Genes Dev 19: 1376–1389

Ding DQ, Sakurai N, Katou Y, Itoh T, Shirahige K, Haraguchi T, Hiraoka Y (2006) Meiotic cohesins modulate chromosome compaction during meiotic prophase in fission yeast. J Cell Biol 174: 499–508

Ding DQ, Matsuda A, Okamasa K, Nagahama Y, Haraguchi T, Hiraoka Y (2016) Meiotic cohesin-based chromosome structure is essential for homologous chromosome pairing in *Schizosaccharomyces pombe*. Chromosoma 125: 205–214

Ding X, Xu R, Yu J, Xu T, Zhuang Y, Han M (2007) SUN1 is required for telomere attachment to nuclear envelope and gametogenesis in mice. Dev Cell 12: 863–872

Drabent B, Bode C, Bramlage B, Doenecke D (1996) Expression of the mouse testicular histone gene H1t during spermatogenesis. Histochem Cell Biol 106: 247–251

Eijpe M, Heyting C, Gross B, Jessberger R (2000) Association of mammalian SMC1 and SMC3 proteins with meiotic chromosomes and synaptonemal complexes. J Cell Sci 113: 673–682

Fraune J, Schramm S, Alsheimer M, Benavente R (2012) The mammalian synaptonemal complex: protein components, assembly and role in meiotic recombination. Exp Cell Res 318: 1340–1346

Fukuda T, Höög C (2010) The mouse cohesin-associated protein PDS5 is expressed in testicular cells and is associated with the meiotic chromosome axes. Genes 1: 484–494

Fukuda T, Fukuda N, Agostinho A, Hernández-Hernández A, Kouznetsova A, Höög C (2014) STAG3-mediated stabilization of REC8 cohesin complexes promotes chromosome synapsis during meiosis. EMBO J 33: 1243–1255

Gómez R, Valdeolmillos A, Parra MT, Viera A, Carreiro C, Roncal F, Rufas JS, Barbero JL, Suja JA (2007) Mammalian SGO2 appears at the inner centromere domain and redistributes depending on tension across centromeres during meiosis II and mitosis. EMBO Rep 8: 173–180

Gómez R, Felipe-Medina N, Ruiz-Torres M, Berenguer I, Viera A, Pérez S, Barbero JL, Llano E, Fukuda T, Alsheimer M et al (2016) Sororin loads to the synaptonemal complex central region independently of meiotic cohesin complexes. EMBO Rep 17: 695–707

Grabarz A, Barascu A, Guirouilh-Barbat J, Lopez BS (2012) Initiation of DNA double strand break repair: signaling and single-stranded resection dictate the choice between homologous recombination, non-homologous end-joining and alternative end-joining. Am J Cancer Res 2: 249–268

Griswold MD (2016) Spermatogenesis: The Commitment to Meiosis. Physiol Rev 96: 1–17

Herrán Y, Gutiérrez-Caballero C, Sánchez-Martín M, Hernández T, Viera A, Barbero JL, de Álava E, de Rooij DG, Suja JA, Llano E et al (2011) The cohesin subunit RAD21L functions in meiotic synapsis and exhibits sexual dimorphism in fertility. EMBO J 30: 3091–3105

Holzmann J, Politi AZ, Nagasaka K, Hantsche-Grininger M, Walther N, Koch B, Fuchs J, Dürnberger G, Tang W, Ladurner R et al (2019) Absolute quantification of cohesin, CTCF and their regulators in human cells. Elife 8: e46269

Hopkins J, Hwang G, Jacob J, Sapp N, Bedigian R, Oka K, Overbeek P, Murray S, Jordan PW (2014) Meiosis-specific cohesin component, *Stag3* is essential for maintaining centromere chromatid cohesion, and required for DNA repair and synapsis between homologous chromosomes. PLoS Genet 10: e1004413

Inselman A, Eaker S, Handel MA (2003) Temporal expression of cell cycle-related proteins during spermatogenesis: establishing a timeline for onset of the meiotic divisions. Cytogenet Genome Res 103: 277–284

Ishiguro KI (2019) The cohesin complex in mammalian meiosis. Genes Cells 24: 6–30

Jin H, Guacci V, Yu HG (2009) Pds5 is required for homologue pairing and inhibits synapsis of sister chromatids during yeast meiosis. J Cell Biol 186: 713–725

Jordan PW, Eyster C, Chen J, Pezza RJ, Rankin S (2017) Sororin is enriched at the central region of synapsed meiotic chromosomes. Chromosome Res 25: 115–128

Kleckner N, Storlazzi A, Zickler D (2003) Coordinate variation in meiotic pachytene SC length and total crossover/chiasma frequency under conditions of constant DNA length. Trends Genet 19: 623–628

Krasikova A, Barbero JL, Gaginskaya E (2005) Cohesion proteins are present in centromere protein bodies associated with avian lampbrush chromosomes. Chromosome Res 13: 675–685

Lee J, Iwai T, Yokota T, Yamashita M (2003) Temporally and spatially selective loss of Rec8 protein from meiotic chromosomes during mammalian meiosis. J Cell Sci 116: 2781–2790

Llano E, Gomez-H L, García-Tuñón I, Sánchez-Martín M, Caburet S, Barbero JL, Schimenti JC, Veitia RA, Pendás AM (2014) STAG3 is a strong candidate gene for male infertility. Hum Mol Genet 23: 3421–3431

Losada A, Yokochi T, Hirano T (2005) Functional contribution of Pds5 to cohesin mediated cohesion in human cells and *Xenopus* egg extracts. J Cell Sci 118: 2133–2141

Mahadevaiah SK, Turner JM, Baudat F, Rogakou EP, de Boer P, Blanco-Rodríguez J, Jasin M, Keeney S, Bonner WM, Burgoyne PS (2001) Recombinational DNA double-strand breaks in mice precede synapsis. Nat Genet 27: 271–276

Moens PB, Marcon E, Shore JS, Kochakpour N, Spyropoulos B (2007) Initiation and resolution of interhomolog connections: crossover and non-crossover sites along mouse synaptonemal complexes. J Cell Sci 120: 1017–1027

Morales C, Losada A (2018) Establishing and dissolving cohesion during the vertebrate cell cycle. Curr Opin Cell Biol 52: 51–57

Muller H, Scolari VF, Agier N, Piazza A, Thierry A, Mercy G, Descorps-Declere S, Lazar-Stefanita L, Espeli O, Llorente B et al (2018) Characterizing meiotic chromosomes’ structure and pairing using a designer sequence optimized for Hi-C. Mol Syst Biol 14: e8293

Nishiyama T, Ladurner R, Schmitz J, Kreidl E, Schleiffer A, Bhaskara V, Bando M, Shirahige K, Hyman AA, Mechtler K et al (2010) Sororin mediates sister chromatid cohesion by antagonizing Wapl. Cell 143: 737–749

Novak I, Wang H, Revenkova E, Jessberger R, Scherthan H, Höög C (2008) Cohesin Smc1β determines meiotic chromatin axis loop organization. J Cell Biol 180: 83–90

Ouyang Z, Yu H (2017) Releasing the cohesin ring: A rigid scaffold model for opening the DNA exit gate by Pds5 and Wapl. Bioessays 39: 1600207

Ouyang Z, Zheng G, Tomchick DR, Luo X, Yu H (2016) Structural basis and IP6 requirement for Pds5-dependent cohesin dynamics. Mol Cell 62: 248–259

Page J, Suja JA, Santos JL, Rufas JS (1998) Squash procedure for protein immunolocalisation in meiotic cells. Chromosome Res 6: 639–642

Parra MT, Page J, Yen TJ, He D, Valdeolmillos A, Rufas JS, Suja JA (2002) Expression and behaviour of CENP-E at kinetochores during mouse spermatogenesis. Chromosoma 111: 53–61

Parra MT, Viera A, Gómez R, Page J, Benavente R, Santos JL, Rufas JS, Suja JA (2004) Involvement of the cohesin Rad21 and SCP3 in monopolar attachment of sister kinetochores during mouse meiosis I. J Cell Sci 117: 1221–1234

Peters AH, Plug AW, van Vugt MJ, de Boer P (1997) A drying-down technique for the spreading of mammalian meiocytes from the male and female germline. Chromosome Res 5: 66–68

Prieto I, Suja JA, Pezzi N, Kremer L, Martínez-A C, Rufas JS, Barbero JL (2001) Mammalian STAG3 is a cohesin specific to sister chromatid arms in meisosis I. Nat Cell Biol 3: 761–766

Prieto I, Pezzi N, Buesa JM, Kremer L, Barthelemy I, Carreiro C, Roncal F, Martínez A, Gómez L, Fernández R et al (2002) STAG2 and Rad21 mammalian mitotic cohesins are implicated in meiosis. EMBO Rep 3: 543–550

Prieto I, Tease C, Pezzi N, Buesa JM, Ortega S, Kremer L, Martínez A, Martínez-A C, Hultén MA, Barbero JL (2004) Cohesin component dynamics during meiotic prophase I in mammalian oocytes. Chromosome Res 12: 197–213

Rankin S (2015) Complex elaboration: making sense of meiotic cohesin dynamics. FEBS J 282: 2426–2443

Remeseiro S, Cuadrado A, Carretero M, Martínez P, Drosopoulos WC, Cañamero M, Schildkraut CL, Blasco MA, Losada A (2012) Cohesin-SA1 deficiency drives aneuploidy and tumourigenesis in mice due to impaired replication of telomeres. EMBO J 31: 2076–2089

Revenkova E, Eijpe M, Heyting C, Hodges CA, Hunt PA, Liebe B, Scherthan H, Jessberger R (2004) Cohesin SMC1 beta is required for meiotic chromosome dynamics, sister chromatid cohesion and DNA recombination. Nat Cell Biol 6: 555–562

Schalbetter SA, Geoffrey Fudenberg G, Baxter J, Pollard KS, Neale MJ (2019) Principles of meiotic chromosome assembly. https://www.biorxiv.org/content/10.1101/442038v2.abstract

Schramm S, Fraune J, Naumann R, Hernandez-Hernandez A, Höög C, Cooke HJ, Alsheimer M, Benavente R (2011) A novel mouse synaptonemal complex protein is essential for loading of central element proteins, recombination, and fertility. PLoS Genet 7: e1002088

Sfeir A, Kosiyatrakul ST, Hockemeyer D, MacRae SL, Karlseder J, Schildkraut CL, de Lange T (2009) Mammalian telomeres resemble fragile sites and require TRF1 for efficient replication. Cell 138: 90–103

Shibuya H, Watanabe Y (2014) The meiosis-specific modification of mammalian telomeres. Cell Cycle 13: 2024–2028

Shintomi K, Hirano T (2009) Releasing cohesin from chromosome arms in early mitosis: Opposing actions of Wapl-Pds5 and Sgo1. Genes Dev 23: 2224–2236

Suja JA, Barbero JL (2009) Cohesin complexes and sister chromatid cohesion in mammalian meiosis. Gen Dyn 5: 94–116

Tedeschi A, Wutz G, Huet S, Jaritz M, Wuensche A, Schirghuber E, Davidson IF, Tang W, Cisneros DA, Bhaskara V et al (2013) Wapl is an essential regulator of chromatin structure and chromosome segregation. Nature 501: 564–568

van Heemst D, James F, Poggeler S, Berteaux-Lecellier V, Zickler D (1999) Spo76p is a conserved chromosome morphogenesis protein that links the mitotic and meiotic programs. Cell 98: 261–271

Viera A, Parra MT, Page J, Santos JL, Rufas JS, Suja JA (2003) Dynamic relocation of telomere complexes in mouse meiotic chromosomes. Chromosome Res 11: 797–807

Visnes T, Giordano F, Kuznetsova A, Suja JA, Lander AD, Calof AL, Ström L (2014) Localisation of the SMC loading complex Nipbl/Mau2 during mammalian meiotic prophase I. Chromosoma 123: 239–252

Wang F, Yoder J, Antoshechkin I, Han M (2003) *Caenorhabditis elegans* EVL-14/PDS-5 and SCC-3 are essential for sister chromatid cohesion in meiosis and mitosis. Mol Cell Biol 23: 7698–7707

Ward A, Hopkins J, Mckay M, Murray S, Jordan PW (2016) Genetic interactions between the meiosis-specific cohesin components, STAG3, REC8, and RAD21L. G3 (Bethesda) 6: 1713–1724

Winters T, McNicoll F, Jessberger R (2014) Meiotic cohesin STAG3 is required for chromosome axis formation and sister chromatid cohesion. EMBO J 33: 1256–1270

Wojtasz L, Daniel K, Roig I, Bolcun-Filas E, Xu H, Boonsanay V, Eckmann CR, Cooke HJ, Jasin M, Keeney S et al (2009) Mouse HORMAD1 and HORMAD2, two conserved meiotic chromosomal proteins, are depleted from synapsed chromosome axes with the help of TRIP13 AAA-ATPase. PLoS Genet 5: e1000702

Wutz G, Várnai C, Nagasaka K, Cisneros DA, Stocsits RR, Tang W, Schoenfelder S, Jessberger G, Muhar M, Hossain MJ et al (2017) Topologically associating domains and chromatin loops depend on cohesin and are regulated by CTCF, WAPL, and PDS5 proteins. EMBO J 36: 3573–3599

Xu H, Beasley MD, Warren WD, van der Horst GT, McKay MJ (2005) Absence of mouse REC8 cohesin promotes synapsis of sister chromatids in meiosis. Dev Cell 8: 949–961

Zhang J, Håkansson H, Kuroda M, Yuan L (2008) Wapl localization on the synaptonemal complex, a meiosis-specific proteinaceous structure that binds homologous chromosomes, in the female mouse. Reprod Domest Anim 43: 124–126

Zhang Z, Ren Q, Yang H, Conrad MN, Guacci V, Kateneva A., Dresser ME (2005) Budding yeast PDS5 plays an important role in meiosis and is required for sister chromatid cohesion. Mol Microbiol 56: 670–680

Zickler D, Kleckner N (1999) Meiotic chromosomes: integrating structure and function. Annu Rev Genet 33:603–754

